# *miR214* regulates sex determination through *gsdf* in zebrafish

**DOI:** 10.1101/2024.05.01.592007

**Authors:** N. Wittkopp, A.M. de Jesus Domingues, R.F. Ketting

## Abstract

Sex determination is a variable and complex mechanism, yet it can be found all over the plant and animal kingdoms. It creates two morphological different outcomes from one and the same species. Our work demonstrates the involvement of the non-coding RNA *dnm3os*, and its embedded microRNA *miR214* in this process for the teleost *Danio rerio*. First, we find that *miR214* acts through *gsdf* to drive female development. Second, additional alleles of *dnm3os* revealed that this lncRNA can also promote male development through yet unknown mechanisms. Finally, we describe that the sex-determining activities of *dnm3os* display a maternal effect, suggesting that imbalances in this gene-regulatory system can be compensated in a stable manner. Although we cannot fully explain the complexity of the mechanisms we have started to reveal, our work once again highlights the complexity and flexibility of sex determination and identifies miRNA and other non-coding RNA mediated activities in this crucial process.

## Introduction

There are many ways how a species can determine its sex. Simplistically spoken, there are species that mainly control it genetically like humans (genetic sex determination GSD) and species where the environment controls the outcome, like turtles (environmental sex determination ESD) (reviewed in (1)). These environmental factors can include, temperature, water salinity or pH, social cues or more generally spoken, stress (2). Moreover, there are species where sex is determined genetically, but can still be overridden by environmental cues (3–6) or species that are in transition between ESD and GSD (reviewed in (7). Besides variability in sex determination mechanisms, there is also variation between species in how individuals maintain their sex. Some species have two sexes and do not change it (gonochorism), some are synchronous or sequential hermaphrodites, some employ one sex (8), and there are mixed forms of the above (2). Clearly, sex determination is a very complex process and gene regulatory mechanisms that affect these have only partially been unravelled.

Specifically in fishes, sex determination is complex. In a BBC Earth documentary in 2017 it was aptly described as ”*Of all the animals, fish are sexually the most fluid*“ (9). Even though the laboratory model organism *Danio rerio* (zebrafish) does not display all possible variations in sex determination, it does possess a rather dynamic system: In wild zebrafish isolates, sex determination seems to be mainly determined by a female heterogametic system: A locus on Chromosome 4 will result in mainly females when present in one copy (10). However, in commonly used zebrafish lab strains, like AB or TUE, this determination system does not seem to be in place anymore which led Wilson et al. in 2014 to the conclusion: Loss of the Natural Sex Determinant in Domesticated Strains (11). Very recently it was proposed, that in zebrafish lab strains, only the female variant of chromosome 4 is present, yet allowing for female and male development (12). Still, this locus on chromosome 4 is differently methylated between males and females in zebrafish lab strains during early sexual differentiation, leading to expression of specific rRNAs in the female (13). But this rather seems to be the consequence of sexual determination and not a cause (14).

What, then, is known about sex determination in zebrafish laboratory strains? During development, zebrafish start as immature females and by the time of 3 to 6 weeks, depending on the growth rate, some larvae undergo metamorphosis: the juvenile oocytes degrade and the male gonad develops (15). This process is influenced by temperature (5, 6), hypoxic conditions (16, 17), rearing density or more generally stress (18). Also, the number of germ cells has a strong effect; without germ cells animals invariably develop into males (19–22). DNA Methylation also seems to play a role in sex determination, as inhibition of methyltransferases prevented the female-to-male transition (23). Along those lines, exposure to higher temperature during embryogenesis led to heat induced males, which displayed enhanced methylation of critical genes involved in sex determination, such as e.g., *sox9a* (24).

In addition to these environmental or physiological effects, different research groups have identified genetic loci that segregate with sex determination in the lab strains, associating *dmrt1* on Chromosome 5 and *cyp21a2* chromosome 16 with sex determination (25), whereas other studies associated chromosomal regions with sex, as the end of chromosome 4 (*sar*) or the middle of chromosome 3 (26, 27).

Despite this wealth of work, the mechanisms driving sex determination are still unknow. The diverse nature of identified factors also points for the existence of other yet unknow regulators. In the present study, we identify a microRNA (miRNA) that affects sex determination. In general, miRNAs are 21-22 nucleotides long RNAs, that do not code for protein. Instead, they operate in a gene-regulatory mechanism that typically dampens the expression of genes by inhibiting translation, accompanied by <u>m</u>essenger <u>RNA</u> (mRNA) destabilization (reviewed in (28, 29)). miRNAs serve as sequence specific guides that bring proteins of the Argonaute family to the 3’ <u>u</u>n<u>t</u>ranslated <u>r</u>egions (3’UTRs) of mRNAs. For effective recognition, only a match of the miRNA seed sequence, formed by nucleotides 2-7 to the UTR, can be sufficient, although additional base-pairing can be required (30). In many cases, miRNAs regulate transcript levels by roughly two-fold, and the effects on translation can be equally modest (31, 32). This implies that these small regulatory RNAs often act in fine-tuning gene expression levels, although switch-like regulation of gene activity has also been described (33). Moreover, the function of a miRNA can be interdependent of additional factors binding to the mRNA’s 3’UTR as for example Brat or Pumilio, which are needed for the destabilisation of the targeted mRNA (34, 35), reviewed in (36).

The miRNA we implicate in sex determination is *miR214*. This miRNA is highly conserved within vertebrates and believed to be derived from one long non-coding RNA that resides in an intron of the *dnm3* gene. This long-non-coding RNA is transcribed opposite to the *dnm3* gene and is hence called *dnm3os* (37, 38). It gives rise to *miR214* but is also home to one of the four *miR199* copies that can be found in zebrafish, *miR199-3*. Unlike *miR199*, the zebrafish genome hosts only one copy of *miR214* and aside from some teleost species, most vertebrates investigated retain only one copy of *miR214* (39). On the opposite strand within an intron of the *dnm3a* coding gene, another miRNA, *miR3120*, is encoded. This so-called ‘mirror-miR’ is annotated in most species found in miRbase, and though it is not annotated in zebrafish in miRBase (40), its presence was confirmed (41).

While a role in sex determination is novel, *miR214* has been described in context of other processes. In the mouse, replacement of the *dnm3os* locus by a LacZ cassette led to aberrant ossification and impaired growth (38). Similarly, human patients carrying a deletion in this gene region display various types of cognitive impairment and skeletal defects (42–45). In mice, an up-regulation of *dnm3os* was found in diabetic mice, where it enhanced inflammation possibly via epigenetic changes through the interaction with Nucleolin (46). One zebrafish study employed synthetic *miR214* and injected it into fertilized zebrafish eggs. They found generally a higher mortality in early embryos and a change in gross morphology including the heart and eye (37). Another study used morpholino antisense oligonucleotides to knock down the expression of *miR214* in early embryos and identified *fused somites* (*sufu*) as a target gene. Here, the slow muscles were affected and one day old embryos displayed an altered somitic structure (47). The same group, using a similar technique, also found *dispatched* 2 (*disp2*) being regulated by *miR214* (48). However, these studies cannot address, whether this phenotype is related to the whole long-non-coding RNA being absent, or the two miRNAs, *miR214* and *miR199* it harbours.

In our study we generated various *miR214* mutant alleles, which revealed a strong effect on sex determination: animals lacking this miRNA more often developed as males, and these males display normal fertility. Using transcriptome analyses, target site predictions and extensive CRISPR-Cas9-mediated mutagenesis we could identify *gsdf* (gonadal somatic cell derived factor) as an important target. *Gsdf* is a teleost-specific gene with a well-established role in sex determination (49). Its modulation by miRNA activity, accompanied by effects on sex determination provides an important new genetic module in zebrafish, and potentially teleost sex determination mechanisms. Additional alleles of the *dnm3os* locus revealed a role in driving male development, and some alleles were neutral in sex-determination. Finally, we uncovered an intriguing maternal effect that seems to enable the animals to restore sex-ratios to normal, even in mutants. Our work establishes *dnm3os* and *miR214*-mediated regulation of zebrafish sex determination, with *gsdf* as an important target. It also highlights the complexity of sex determination in general, and the *dnm3os/ miR214* locus in particular.

## Results

### The *dnm3os* locus

*Dnm3os* is a non-coding RNA harbouring *miR199-3* and *miR214* (Fig 1A). Its expression in mouse is reported to be in the mesodermal lineage, mainly in the bone forming tissue (38). In zebrafish we could recapitulate its expression by creating a transgenic line with the *dnm3os* promotor driving a fusion of GFP and the non-coding *dnm3os* RNA: *Tg(dnm3os:GFP-dnm3os)* (Fig S1C). Therefore, this RNA appears to be highly conserved between the mammalian and teleost lineages. Replacement of this locus with a lacZ-reporter-cassette in mouse led to neonatal death, with an early ossification phenotype (38). In human, a heterozygous deletion of this genomic region manifests in mental and skeletal abnormalities (42–45). Based on this evidence, we asked if this phenotype could be reproduced in zebrafish and employed CRISPR-Cas9 (50, 51) to generate different deletion alleles of the *dnm3os* locus, including specific deletions for both miRNAs. To our surprise, neither heterozygous nor homozygous *dnm3os^Δ2.3kb^* mutants (Fig 1A), lacking the complete lncRNA, showed any obvious defects, neither in bone structure nor sex ratios - even when kept homozygous over several generations (Fig S1A, B). A 51bp deletion targeting *miR199-3* specifically also had no detectable phenotype (data not shown). Finally, a deletion of 180bp including the whole *miR214* and spreading 80bp up-stream, *miR214^Δ180bp^* (Fig 1B) also showed no phenotype (Fig 1C graphs 1 & 2). We conclude that loss of large fragments of *dnm3os* lncRNA does not trigger the phenotypes that would be expected based on work in mammals. However, when we specifically deleted *miR214*, we did observe a phenotype.

**Figure 1.**
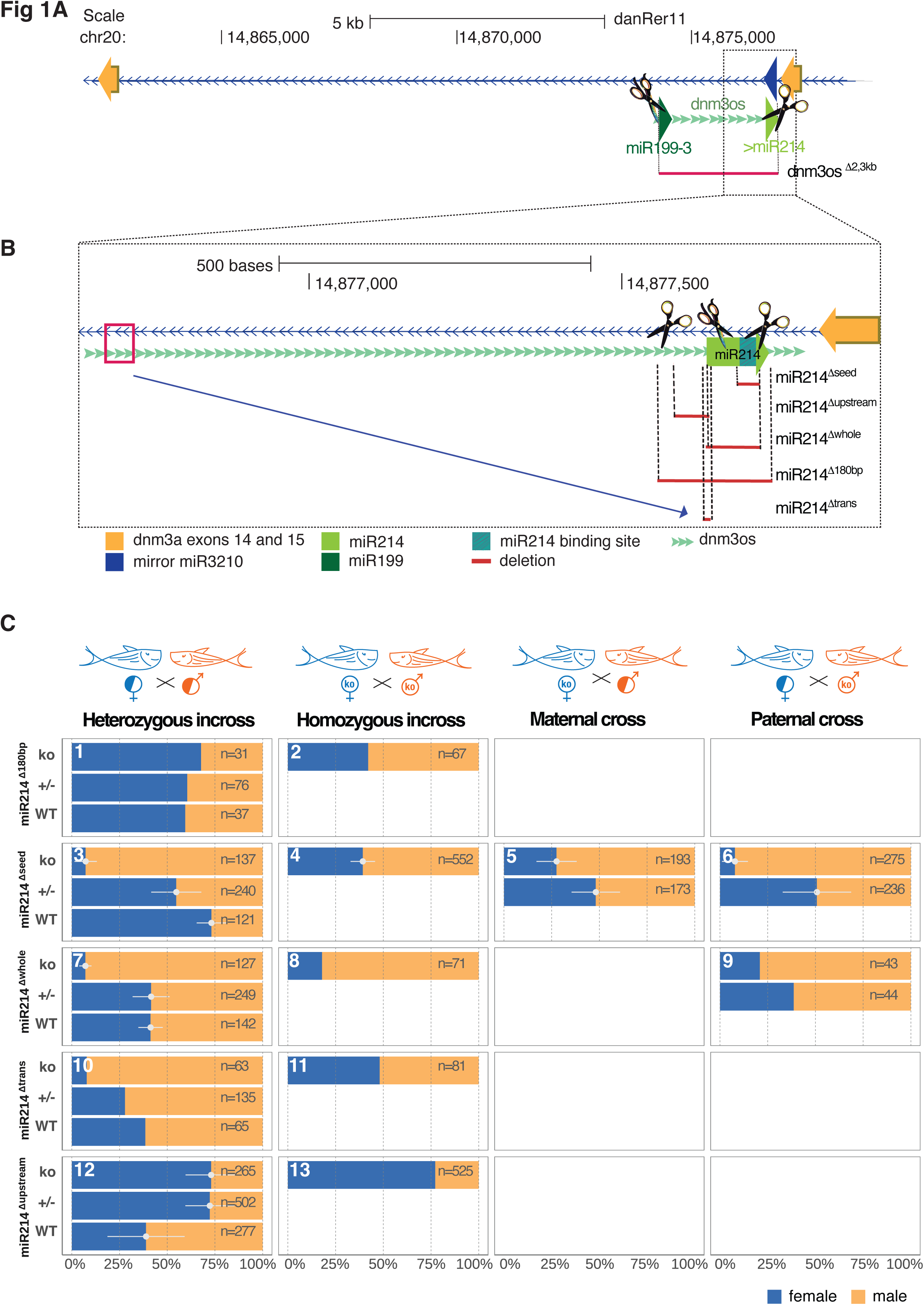
(A): Gene locus encoding for *dnm3os*, which is a precursor long non-coding RNA (lncRNA) harboring the two miRNAs *miR199-3a* and *miR214*. It is located on the antisense strand of an intron spanning the exon 14 to 15 of the coding gene *dnm3a*. The depicted scissors highlight the binding of used CRISPR sgRNAs resulting in the *dnm3os^Δ2.3kb^* mutation. (**B)** is a magnification of the *miR214* encoding part of *dnm3os*, with all different *miR214* mutant alleles. The scissors here also depict the side of different sgRNAs used for mutagenesis. (C) shows different type of crosses for various *miR214* mutant alleles. The type of cross is symbolized by the fish pictograms on top, the y-axis shows the genotype of the offspring of those crosses, the x-axis gives the sex distribution in percentage. The actual number of evaluated fish can be found within the bars. Error bar represents standard deviation between at least three crosses.

### Mutation in *miR214* lead to a male bias

A deletion of 41bp was introduced by one sgRNA and eliminated the whole mature miRNA sequence, including the seed sequence, *miR214^Δseed^* (Fig 1B). Physiologically, these fish did not display any obvious defects. Changes in body length, as reported for the *dnm3os* ko mouse (38), were also not detected (Fig S2A). We noted, however, that we obtained hardly any homozygous females from crosses between heterozygous parents (Fig 1C graph 3). Typically, less than 10% of the homozygous mutant offspring developed into females, whereas their siblings, which were raised in the same tanks and hence under identical conditions, did not display this effect.

To substantiate our observation on a molecular level, we performed <u>q</u>uantitative <u>R</u>everse <u>T</u>ranscription PCR (qRT-PCR) on isolated one month old gonads, a timepoint when sex differentiation has started, but is not yet visible. We reared all siblings from a cross between heterozygous females with homozygous males together, and then dissected the gonads, employing the *Tg(vasa:eGFP)* line , which marks germ cells (52). Non-germline tissue was used to assess the genotype of each individual and up to five gonads were pooled. Then, we interrogated *dmrt1* using qRT-PCR, a known determinant for testis development in zebrafish (53). *Dmrt1* showed an at least 1.5-fold increase in *mir214* mutants, indicating that sex determination is male-biased already in one month old *miR214^Δseed/Δseed^* larvae (Fig S2C).

### Loss of *miR214* does not result in a loss of primordial germ cells

As the number of germ cells is known to have a strong influence on sex determination, with loss of germ cells leading to male development (19–22), we counted the number of germ cells, including the number of germ cells that did not reach the gonadal niche in one day old (1dpf) embryos, using again the *Tg(vasa:eGFP)* transgenic line (52) (Fig S2E, F). Even though germ cell counts were lower in mutant compared to wild-type animals, similarly low germ cell counts were found in heterozygous siblings, indicating that a reduction in germ cell number does not explain the sex bias (Fig S2D).

### Male bias in additional *miR214* alleles

Because the *dnm3os^Δ2.3kb^* and *miR214^Δ180bp^* alleles did not show a sex determination phenotype, and because sex determination is so fluid in zebrafish, with different background strains and families having a sex bias on their own (27), we wanted to make sure the reported phenotype of *miR214^Δseed^* did not stem from off-target effects of the used sgRNA and/or from strain-related biases. Therefore, we created further alleles of *miR214* using a different sgRNA in a zebrafish strain that we freshly obtained from the eZRC (*alb^ti9^* in TUE/AB). One allele deleted 87bp, which nearly ablated the whole *miR214* precursor, and was therefore termed *miR214^Δwhole^* (Fig 1B). Another allele is a translocation of 44 nucleotides from upstream of the *dnm3os* locus being inserted into the 5’ part of the mature miRNA, *miR214^Δtrans^* (Fig 1B), rendering it improbable to be processed into the mature *miR214* form. We tested the expression for *miR214* using qRT-PCR at least for two alleles, in *miR214^Δseed^* and *miR214^Δwhole^* in 5dpf embryos, one month old and 6 weeks old (6wpf) gonads (Fig S3A, C, E). In all cases, expression was below detection levels and therefore we assume these alleles represent complete knockouts. Both *miR214^Δwhole^* and *miR214^Δtrans^* resulted in a male bias of homozygous offspring from heterozygous parents (Fig 1C graphs 7 & 10). We conclude that *miR214* regulates sex determination in zebrafish. Specifically, it supports female development as its absence leads to over proportionally many males.

### A female-biased allele of *miR214*

Just upstream of *miR214* we detected a predicted ZNF418 Transcription Factor binding site using JASPAR (54). To probe the possibility that this element may regulate *miR214* expression, we generated a 40bp deletion that hit this upstream element: *miR214^Δupstream^* (Fig 1B). Animals homozygous or heterozygous for this allele displayed a strong female bias compared to wildtype siblings, with around 70% of the animals developing as females (Fig 1C graphs 12 & 13). This would argue for a repressive activity of ZNF418, which is in accordance with published data (55). However, qRT-PCR experiments did not support this simple model. We did not observe any up-regulation of *miR214* in one month old gonads or 5dpf embryos (Fig S3A, C). Rather, we detected down-regulation in both homozygous and heterozygous *miR214^Δupstream^* animals. This was in the same range, as animals heterozygous for *miR214* deletions. To our further surprise, we observed a 4-fold down regulation of *mir214* in 6wpf gonads from homozygous *miR214^Δupstream^* fish, specifically in females, while the male gonad did not show this effect (Fig S3E). We also tried to investigate, if *miR214^Δupstream^* might be dominant over the *miR214^Δseed^* allele, but trans-heterozygous *miR214^Δupstream/Δseed^* animals did not show any sex bias (Fig S4A).

Clearly, this data does not agree with the hypothesis that ZNF418 mediates repression of *miR214*, combined with a general female-fate promoting activity of *miR214*. However, it is possible that this model does apply at a specific developmental time-point, or in a specific cell type, that we have thus far failed to analyse. Possibly, *miR214^Δupstream^* affects other aspects of *dnm3os* besides *miR214*, which may explain a female biased phenotype.

### Identification of potential *miR214* targets

To identify potential *mir214* targets we used two approaches. First, we tested candidate genes using qRT-PCR in one month old gonads from individuals from a cross with a *miR214^Δseed/Δseed^* male to a *miR214^Δseed/+^* female. We would expect that *miR214*-targeted transcripts increase in abundance in *mir214^Δseed/Δseed^* mutants. Our first candidate was *ezh2,* a known target gene for *miR214* in mouse (56), however in our experiments it was rather down-regulated in *miR214^Δseed/Δseed^* mutants (Fig S2C). Additionally, analysis of its 3’UTR did not show any *miR214* binding sites, rendering it an unlikely target in zebrafish. Next, we identified candidates through TargetScanFish (Release 6.2 (57, 58)), a program that predicts miRNA target sites in transcripts. From these predictions we tested *gsdf, nup54, klhl8, mei1 (*ENSDARG00000090972.1*), fam110a* and *ptgdsb*. Three of the predicted targets, namely *nup54, klhl8* and *fam110a* showed no significant expression changes, whereas *mei1* and *gsdf* did. Especially *gsdf* displayed a 3-fold up-regulation in homozygous *miR214^Δseed^* gonads (Fig S2C).

As a second approach to identify potential *miR214* targets, we used RNA-sequencing of whole gonads. We used gonads from one month old animals, a timepoint when sex differentiation begins. We reared offspring from a cross between *miR214^Δseed/Δseed^* mutant fathers and *miR214^Δseed/+^* mothers, isolated the gonads with the aid of the *Tg(vasa:GFP)* transgene (52) and used the somatic part of the animals for genotyping. Using <u>d</u>ifferential <u>g</u>ene <u>e</u>xpression (DGE) we then identified up and downregulated genes. For clarity: animals were not separated according to their sex since this was not yet apparent at the time of analysis. With the specified cut-off of p_adj_<0.05, we found a total of 3208 genes up-regulated and 2974 down-regulated (Fig 2A). Consistent with the male bias in *mir214* mutants, we observed a tendency of male-expressed genes like *amh* (2.3-fold), *sox9a* (1.7-old) or *star* (2.4-fold), being expressed at a higher level in *mir214^Δseed/Δseed^* gonads compared to heterozygous control gonads (Fig 2C). *Gsdf,* which we had already analysed before using qRT-PCR, could once more be identified as a gene that is upregulated in *mir214^Δseed^* mutants. Furthermore, we identified some genes that are involved in the retinoic acid (RA) pathway. *miR214* has already been suggested to play a role in the RA pathway (discussed in (59)), and at least some members of the RA pathway are predicted to have at least one *miR214* binding site and are implied in sex determination: *aldh1a2* (RA synthesis) (60), *rxraa* and *rxrba* (RA receptor) (61, 62), *sox9a* (RA target gene) (63), *wt1a* and *wt1b* (RA target genes) (64), *cyp26b1* and *cyp26c1*(RA degrading enzyme) (60, 65) and *nr5a1* (SF1 ortholog, transcription factor) (66, 67).

**Figure 2:**
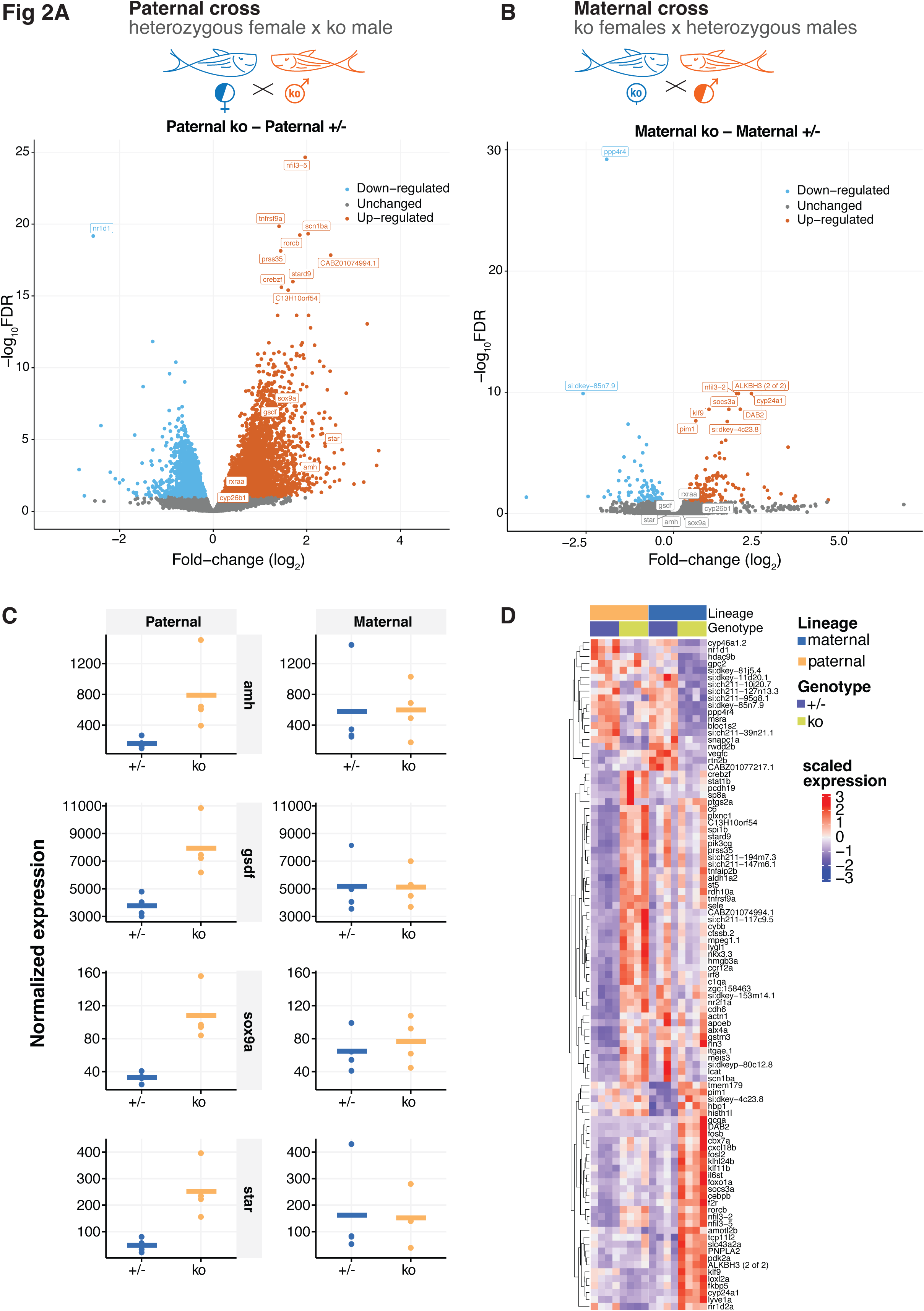
Comparison of RNA expression in one month old gonads between *miR214^Δseed^* homozygous and heterozygous siblings derived either from a paternal cross in (**A)** (homozygous *miR214^Δseed^* males with heterozygous *miR214^Δseed^* females) or a maternal cross in (**B)** (homozygous *miR214^Δseed^* females with heterozygous *miR214^Δseed^* males). Expression of genes associated with testis development are highlighted in (**C**), which showed differential expression between homozygous and heterozygous animals derived from a paternal cross (homozygous *miR214^Δseed^* males with heterozygous *miR214^Δseed^* females) on the left panel site. These genes did not change significantly in larvae derived from a maternal cross (homozygous *miR214^Δseed^* females with heterozygous *miR214^Δseed^* males) on the right site. The 50 most differentially regulated genes were compared in a heat map between the paternal and maternal cross in (**D)**.

### 3’UTR alleles of *gsdf* induce a male bias

To probe the relevance of these target genes for the sex determination phenotype we generated mutants for *gsdf* and the RA-pathway components *cyp26b1* and *rxraa*, by deleting the binding sites for *miR214* in their 3’UTRs. For the components of the RA acid pathway (Fig S5A, B), we did not observe a clear sex bias in the homozygous mutants compared to their heterozygous or WT siblings, neither in single mutants (Fig S5C, D), nor in double mutants (Fig S5E).

*Gsdf* is a known initiator of male sex in Medaka (*Oryzias latipes*), where its overexpression led to sex reversal of genetic females into phenotypic males. Conversely, loss of *gsdf* triggered a male to female transition (68–70). In zebrafish, however, a lack of *gsdf* did not alter sex ratios, but rendered homozygous females sterile over time (71). A gain of *gsdf* function, which would be the result of loss of miRNA action on *gsdf*, has not been studied in zebrafish thus far. Zebrafish *gsdf* harbours three potential *miR214* binding sites in its 3’UTR (Fig 3A), one of which is conserved in some teleosts (Fig S2B). Using two sgRNAs we obtained four *gsdf* alleles with lesions in the 3’UTR (Fig 3A). All of these alleles deleted the conserved *miR214* binding site in the middle of the 3’UTR, which was predicted by TargetScan (57, 58). The other two potential *miR214* sites further upstream in the 3’UTR were not affected in these alleles. Two of these alleles did not display any sex distribution alterations when homozygous: *gsdf^Δ100bp^* and *gsdf^Δ200bp^* (Fig 3B, graphs 1 - 4). In contrast, the *gsdf^Δ400bp/Δ400bp^* and *gsdf^Δ800bp/Δ800bp^* alleles, which shared an additional 200bp deletion upstream of the conserved *miR214* site, copied the *miR214^Δseed^* ko phenotype (Fig 3B, graphs 5 & 7): only around 10% of the animals were female, and *gsdf* mRNA levels were upregulated 4-fold in *gsdf^Δ400bp/Δ400bp^* animals (Fig S3D). We then incrossed *gsdf^Δ400bp/+^, miR214^Δupstream/+^* double heterozygous fish to probe which of the alleles would be dominant over the other. We found that the homozygous *gsdf^Δ400bp/Δ400bp^* offspring were mainly males irrespective of *miR214* status, and the fish which carried *miR214^Δupstream^* had the tendency to be female, but only if the *gsdf* locus was wild-type (Fig S4B). Therefore, the *gsdf^Δ400bp^* is dominant over the *miR214^Δupstream^* allele, consistent with the idea that *gsdf* acts downstream of *miR214/dnm3os*.

**Figure 3.**
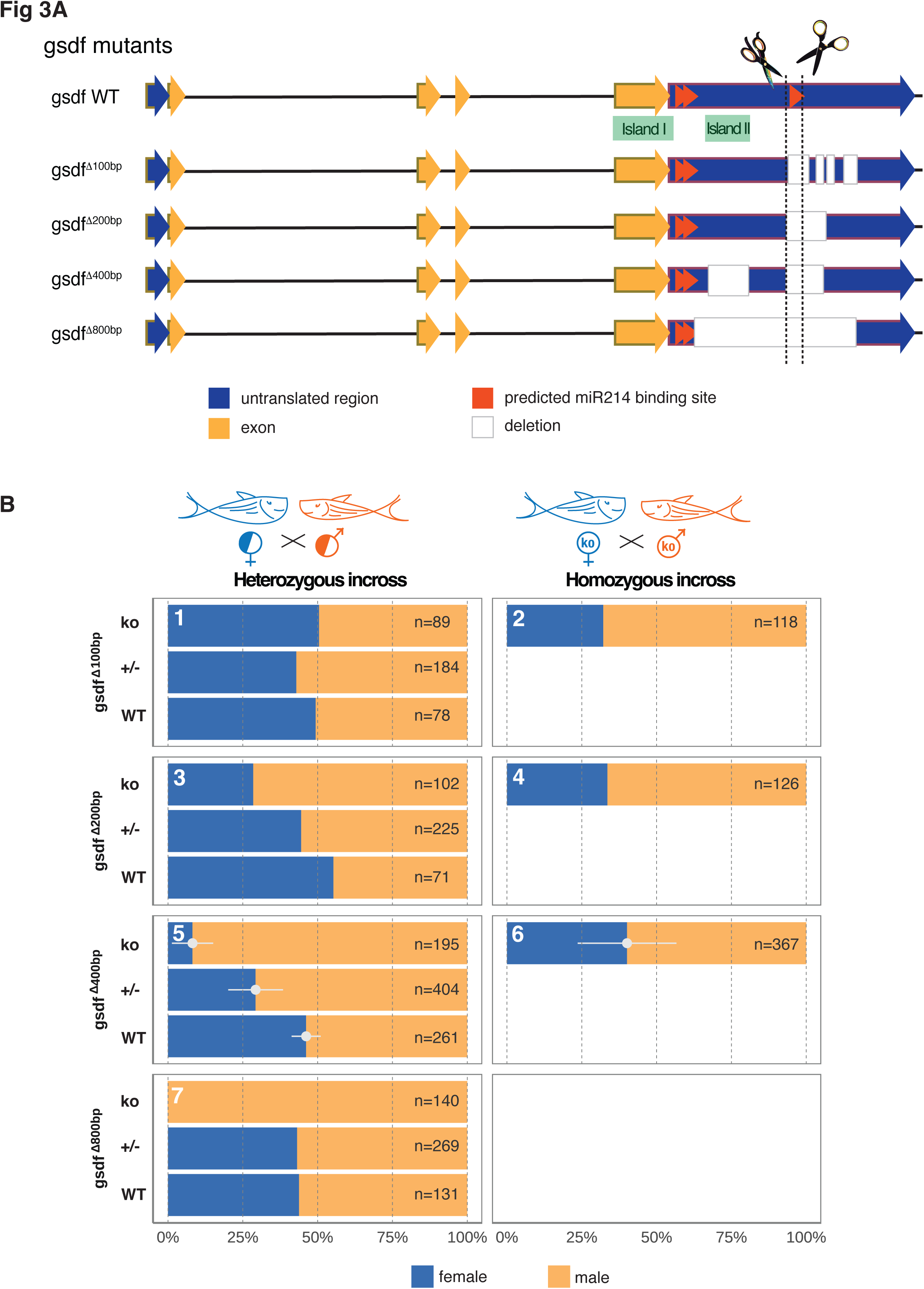
(A): Different alleles generated for *gsdf*, ordered according to their size of deletion. Scissors mark the sequence where the sgRNAs cut and red arrows highlight potential *miR214* binding site. Note that only the last one is evolutionary conserved and predicted by TargetScanFish (57, 58). In **(B)** the sex distribution of the different alleles was assessed, where the allele and the genotype are on the y-axis and the distribution in percentage on the x-axis. In case of at least three different crosses the standard deviation is added. The genotype of the parents is depicted on top with the different fish cartoons.

We conclude that *gsdf* is a feasible sex-determination-relevant target of *miR214*. However, other regulatory elements are likely present within the 3’UTR of *gsdf*, and loss of one conserved *miR214* target site is not sufficient to de-regulate *gsdf* sufficiently to trigger a sex-bias phenotype.

### A maternal effect on sex determination

The homozygous mutant offspring from heterozygous incrosses, as well as from homozygous mutant fathers, always displayed a strong male bias as described above. This holds for the sex-biased *miR214* (Fig 1C, graphs 3, 6, 7, 10), as well as for the *gsdf* alleles (Fig 3B, graphs 5 & 7). We were astonished to find, however, that sex ratios were nearly restored to wildtype levels in homozygous mutant offspring of homozygous *miR214^Δseed^* females (Fig 1C graph 4). Similarly, when crossing females homozygous for the *miR214^Δtrans^* (Figa 1C graph 4&5) and *gsdf^Δ400bp^* alleles (Fig 3B graph 6), sex ratios were close to wildtype in the homozygous mutant offspring, albeit with a rather broad range of fluctuations for *gsdf^Δ400bp^*. Accordingly we could observe in one month old gonads, that *gsdf* level were going back to normal in offspring of homozygous mutants, compared to offspring of heterozygous parents (Fig S3D). An exception was the *miR214^Δwhole^* allele: homozygous mutant offspring of homozygous *miR214^Δwhole^* females still displayed a male bias (Fig 1C graph 8). Likewise, the *miR214^Δupstream^* allele did not show a maternal effect (Fig 1C graph 13): homozygous mutants displayed a strong female bias of ∼70% irrespective of whether the mother was homozygous or heterozygous (Fig 1C graph 12 & 13).

We also looked at this maternal effect on a molecular level, using mRNA-sequencing. As already described, in homozygous *miR214^Δseed^* mutants derived from heterozygous mothers, we identified 3208 genes up-regulated and 2974 down-regulated (cut-off padj<0.05) between homozygous and heterozygous siblings (Fig 2A and supplemental table 1). In contrast, when we compared homozygous to heterozygous siblings from homozygous *miR214^Δseed^* mothers, we only detected 55 upregulated genes and 101 downregulated genes in the *miR214^Δseed/Δseed^* animals (cut off padj<0.05) (Fig 2B and supplemental table 2).

We compared the top most differentially expressed genes, 50 for each of the comparisons, to observe if there were any patterns. Specifically, we expected that up-regulated genes in the *miR214^Δseed/Δseed^* offspring of heterozygous mothers would not be differentially expressed in the *miR214 ^Δseed/Δseed^* offspring of homozygous mothers. With few exceptions, this was mostly true (Fig 2D). We conclude that a maternal effect arises in the gonads of *miR214* mutant females that can compensate for the loss of *miR214* in their offspring. These results also show that the male bias is not due to secondary mutations that linger in the background of the mutants we generated.

### Molecular analysis of the maternal effect

We aimed to understand how the sex bias can disappear in offspring of homozygous *miR214^Δseed^* mothers. We reasoned that the mis-regulation of *gsdf* in these females may trigger compensatory effects on gene regulation, which may then in turn be inherited to the offspring. As inheritance of gene activity state is often coupled to DNA methylation, we probed the DNA methylation status of CpG islands in the following genes: *gsdf* (Fig S6a), *sox9a* and *cyp26b1* (Fig S6B, C). We also included a region of the fem-rDNA region (Fig S6D), which is de-methylated during gonad development only in female zebrafish (13). For each of these genes we used methylation-specific PCR to assess the methylation status of annotated CpG islands in 1 month old gonads from wildtypes, or in homozygous mutants that were derived from either homozygous *miR214^Δseed^* mutant fathers or mothers. We employed two different primer-sets, hoping to avoid sequence-specific effects. We did not find any DNA methylation differences for *sox9* (Fig S6B), *cyp26b1* (Fig S6C) and fem-rDNA (Fig S6D). However, we detected potential hypermethylation at the *gsdf* locus, specifically in homozygous mutant offspring of homozygous *miR214^Δseed/Δseed^* mutant mothers, suggesting it is hyper-methylated (Fig S6A).

Next, we examined the *gsdf* locus in more detail and probed two CpG islands, which we will refer to as island one (Is1) and island two (Is2) (Fig S7A), by bisulfite-sequencing. Is1 is located in *gsdf’s* last exon stretching over its C-terminus, Is2 is situated in its 3’UTR overlapping with the mutations *gsdf^Δ400bp^* and *gsdf^Δ800bp^* (Fig S8A), the two *gsdf* alleles that display a sex bias (Figa 3 B graphs 5 & 7). We assessed the methylation status of these two islands in one month old gonads of *miR214^Δseed^* (Fig S7B, D), *miR214^Δwhole^* (Fig S7C, E) and *gsdf^Δ400bp^* mutants (Fig S8B). As mentioned before, the *miR214^Δseed^* mutation loses its sex bias in individuals derived from homozygous mutant mothers (Fig 1C graphs 4 & 5), as does *gsdf^Δ400bp^* (Fig 3B graph 6), albeit with more variation between crosses. By contrast, the *miR214^Δwhole^* allele always displays a male bias, regardless of the parents’ genetic make-up (Fig 1C, graphs 7, 8, 9).

DNA methylation in Is1 was generally high and was not affected by the genotype of the mother (Fig S7B, C). Is2, however, showed interesting dynamics: DNA methylation appeared to be higher only in the mutants derived from a *miR214^Δseed^* homozygous incross, and not in offspring from a heterozygous cross (Fig S7D). This effect was not seen for *miR214^Δwhole^* (Fig S7E), and *gsdf^Δ400bp^* could not be analysed for that region (Fig S8A) and thus cannot explain the compensatory effect in *gsdf^Δ400bp/Δ400bp^* mutants from homozygous mothers. Even if we cannot demonstrate causality, these results are consistent with the idea that upregulation of *gsdf* at the post-transcriptional level may be countered by repressive effects at the transcriptional or chromatin levels.

## Discussion

We propose a model where sex determination in zebrafish is modulated by miRNA action; specifically, *miR214* regulates *gsdf* expression to allow for male and female development (Fig 4B). If this regulation is disturbed by either *miR214* itself being absent or by *gsdf* 3’UTR mutations, this leads to an up-regulation of *gsdf*, counteracting female development (Fig 4C). However, if homozygous animals do develop as female, *gsdf* expression is again reduced to wildtype level in her offspring, irrespective of their genotype, and sex ratios are balanced once again (Fig 4D, right site). This implies that there is a feedback mechanism, which we so far could not identify. Interestingly, this compensatory effect is not seen in all mutants, as *miR214^Δwhole^* mutants keep the male bias (Fig 4D, left site) and *miR214^Δupstream^* mutants a female bias (Fig 4A). Finally, not all alleles of the miRNA and *gsdf* loci produce the expected phenotypes, indicating further genetic complexity. Besides a recent paper showing that loss of *miR-17∼92* leads to a complete male to female sex-reversal in mice (72), to our knowledge, our work describes only the second case of a miRNA involved in sex determination in a vertebrate. Below, we will go deeper into the role of *miR214* and into the complexities we uncovered with our experiments.

**Figure 4:**
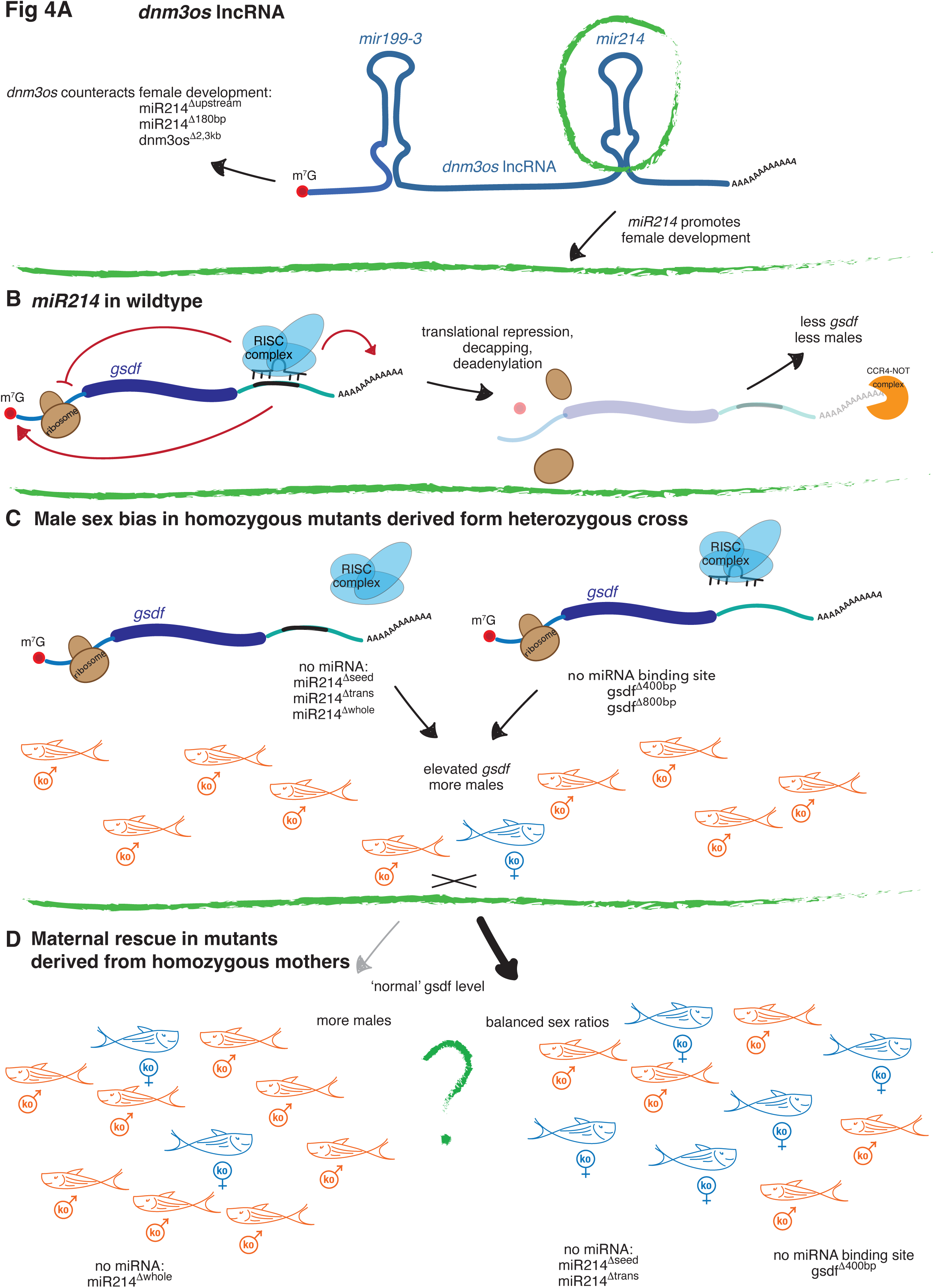
The whole lncRNA *dnm3os* might have different roles in sex determination (**A**). Whereas *dnm3os* counteracts female development, especially *miR214* promotes the female fate. In a wildtype scenario *miR214* would be bound by the RISC and *gsdf* would be targeted by it, leading to reduced mRNA levels of *gsdf* by translational repression, decapping and/or deadenylation **(B)**. In case of a *miR214* mutation (**C**, left side) the RISC cannot be guided to *gsdf* or any other target mRNA and thus the target mRNA is stabilized. Similarly, *gsdf* will be stabilized if the *miR214* binding site is removed as in the *gsdf^Δ400bp^* and *gsdf^Δ800bp^* mutants. In both scenarios, the homozygous offspring of incrossing heterozygous carriers, does show a male bias. If then again homozygous carries are incrossed or a homozygous female is crossed to a heterozygous male (**D,** left site), the homozygous offspring loses this male bias in most cases, an exception is the *miR214^Δwhole^* allele, where this effect seems to be, if all, weak (**D**, right site).

### miRNA mediated sex determination

Although *miR214* has mainly been studied in the context of bone development, there is pre-existing literature about *miR214* in the gonad. For example, *miR214* was identified in an expression analysis between male and female zebrafish gonads, as being altered by templated addition of nucleotides, which might affect its ability to target and regulated mRNAs (73). Moreover, it was identified in ovaries from Atlantic Halibut (*Hippoglossus hippoglossus*) (74), the pufferfish *Takifugu rubripes* (75), pigeon (76) and in the follicular fluid of bovine ovary (77). It was also found to be de-regulated in human malignant ovarian germ cell tumors (78–80) and in mouse, *miR214* has been implicated in the entry of meiosis during sexual differentiation, mediated via the retinoic acid pathway (59).

Interestingly, *miR214* seems to also play a role in fat cell metabolism in mice and humans (81) as well as ducks (82). According to our data *miR214* influences the expression of *gsdf*, which in turn is known to have an impact on the zebrafish’s fat and insulin metabolism (83). A connection between the energy metabolism and fertility are well known and seem to be preserved over evolution (reviewed in (84, 85)). This link also exists in humans, as during the female menstruation cycle the body changes its susceptibility to insulin (86, 87). Moreover, the lncRNA *dnm3os*, which harbours *miR214* and *miR-199-3a*, was shown to be up-regulated in response to diabetes in rats and humans (46). Overall, it transpires that *dnm3os* and its miRNAs may affect the physiology of animals in various ways, and it seems likely that a clean dissection of individual regulatory units will be hard to achieve.

### Post-transcriptional control of *gsdf*

We identified *gsdf* as a likely target of *miR214*. However, we have to note that deletion of just the *miR214* binding site in the 3’UTR did not phenocopy loss of *miR214*. Only a larger deletion of the 3’UTR did, suggesting that there are additional factors that affect *gsdf* expression. One may even argue that the effect of the *gsdf* 3’UTR mutation does not involve *miR214* at all. However, a double mutant between the hypermorphic *miR214^Δupstream^* allele and *gsdf^Δ400^* allele showed the *gsdf^Δ400^* phenotype, consistent with a model in with the *miR214^Δupstream^* allele acts through the deleted *miR214* binding site in the *gsdf* 3’UTR.

While we cannot cleanly isolate the effect of *miR214* from other factors that act on the *gsdf* mRNA, it is clear that the *gsdf* 3’UTR provides important regulatory information. Next to miRNA binding sites, several binding sites for Pumilio can be found (RBPDB database (http://rbpdb.ccbr.utoronto.ca/) (88)), suggesting that Pumilio, and likely additional RNA-binding proteins bind this 3’UTR. Possibly, these factors facilitate the binding of *miR214* to its target, as has been shown before for other miRNA-target pairs (34, 35), or otherwise affect *gsdf* mRNA activity and or stability. Our work cannot further dissect these mechanisms but does show that *gsdf* activity is regulated through several post-transcriptional mechanisms acting through its 3’UTR.

### Compensatory effects

Sex determining genes have evolved several times in evolution. A theory after Wilkins states that there is an optimal range of sex ratios for any given population. If the sex ratio is skewed by a mutation, new mutations will be favoured that drive the ratio back to its optimum and by this creating new sex regulatory loops (89, 90). This theory might be too passive, as the organism must remain competitive to ensure species survival. Therefore, there may be mechanisms in place that counteract changes in sex bias within a population faster and more effectively. Such effects may work epigenetically, as shown in the plant *Arabidopsis thaliana* (91). Alternatively, buffering mechanisms may exist, such as genetic compensation, which was first reported in zebrafish, when a morpholino knockdown displayed a phenotype, but the corresponding mutation did not (92). We found that the sex bias phenotype of many of the generated alleles could be compensated in those animals that did develop into females, even though the population experienced a strong male bias. How can this work?

We detected subtle hyper-methylation of the *gsdf* locus as a potential response: such hypermethylation was only detected in gonads of offspring from homozygous mutant females. Therefore, it is possible that this hypermethylation reflects transcriptional downregulation of *gsdf*. We note that the affected CpG island overlaps at the genomic level with the region coding for the 3’UTR of *gsdf*, and in particular the region we identified to affect sex ratios. This further complicates the interpretation of the alleles that affect the 3’UTR of the *gsdf* gene, as the gene may also be affected at the transcriptional level.

We also looked into the polycomb member *ezh2*, which modifies methylation (93) and is a known *miR214* target in mouse (56), but in zebrafish there is no predicted *miR214* binding site and its expression is not up-regulated in *miR214^Δseed/Δseed^* mutant gonads, as we have tested using qRT-PCR. *Ezh1* might be another candidate and harbours a *miR214* binding site according to TargetScanFish (57, 58), but was also not up-regulated in our mRNA-sequencing data set.

Finally, the *dnm3os* lncRNA itself is suspected to be involved in epigenetic modifications by binding to Nucleolin (46). Its expression is activated via NF-κB binding at the conserved *dnm3os* promotor region (37, 46) under diabetic conditions in rats. This is interesting, as the target gene of *miR214*, *gsdf*, also regulates metabolic processes, as was detected in *gsdf* mutants, which displayed down-regulation of the insulin-receptor and stored excess lipids (83). We can therefore speculate that upon initial overexpression of *gsdf* in *miR214* mutants, feedback mechanisms are activated through metabolic processes that are yet not clearly defined. We expect that a full understanding of the compensatory mechanism(s) that act upon loss of *miR214* will require multi-faceted approaches, including dissection of gene regulatory mechanisms as well as metabolism.

### Differential effects of specific alleles: additional functions of *dnm3os* lncRNA?

We generated different alleles of *dnm3os*, and not all of them displayed a sex determination phenotype, even if they did hit the *miR214* stem-loop. What could be the reason(s) behind this observation? Given that particularly the larger deletions did not display a phenotype, one possibility is that the *dnm3os* lncRNA affects sex determination in both directions (Fig 8A), and that the larger alleles take out both functions, effectively not resulting in a phenotype. It is hard to speculate how such differential effects can be explained without knowing how the *dnm3os* lncRNA acts. Possibly, the 3D-folding of the RNA is involved, and some alleles may disrupt essential folds, or others may take out specific structures. To address these questions a closer investigation of the *dnm3os* lncRNA and its binding partners will be required. We note that, in general, removal of complete lncRNAs often had no phenotype in zebrafish (94), just like we found for *dnm3os*. Nevertheless, we show that smaller deletions can reveal phenotypes. Maybe, tackling the function of lncRNAs through more subtle deletions and manipulations, rather than complete deletions, will proof to be a more fruitful approach in addressing lncRNA function.

### The TGF-beta pathway in sex determination

*Gsdf* acts as a receptor ligand (https://www.uniprot.org/uniprotkb/B0ZE98/entry) in the TGF-beta pathway. It is conserved in teleosts, but is not present in therians, where it was lost. Interestingly, the synteny of the chromosomal fragment/environment, which harboured *gsdf* can still be traced up to humans (95). And although *gsdf* itself is not preserved outside the teleost clade, the TGF-β pathway it belongs to is often found to be involved in sex determination and differentiation with members like *amh*, *amhr2*, *bmpr1b* and *gdf6* (reviewed in (96)). Paralogs of *gsdf*, which all might have arisen during the vertebrate genome duplication are *bmp15* and *gdf9* and are conserved in vertebrates (83). Mutations in both are implicated in ovarian failure in zebrafish (97, 98), but also in human (99, 100). The involvement of the TGF-β pathway in sex determination seems to be an evolutionary conserved theme that can be traced down to invertebrates like the worm *Caenorhabditis elegans* (101) or the fruitfly *Drosophila melanogaster* (102) and is still present in mammals (reviewed in (103, 104)). Therefore, although *gsdf* itself has been lost in mammals, the general mechanisms behind it, possibly including its regulation by *miR214*, might be ancient and conserved from invertebrates to mammals.

## Material and methods

### Primer and sgRNA design

General primer were designed using NCBI’s Primer-Blast function. For methylation specific primers, the MethPrimer Tool was employed (105) and for sgRNA design we used the CRISPR/Cas9 target UCSC tracks provided by (106) and the CRISPRScan Cas9 UCSC tracks from (107).

### Preparation of sgRNA

10µM of each primer were annealed by placing them in boiling water and letting them cool down to 4C overnight. These duplexes were then ligated into BsaI cut DR274 (Addgene 42250, (50)), containing either a T7 or Sp6 promotor, sequenced and cut with DraI. The resulting fragment was gel-purified and then used to be transcribed using either MegaScript kits from Ambion or HiScribe kits from NEB according to manufacturer’s instructions. A DNase digest was added, and the RNA precipitated overnight at -20C. Then concentration and integrity were assessed via gel electrophoreses. For injection into eggs, we used: 20µg sgRNA or more, 0.5 µl Phenol Red solution (Sigma, P0290), 2 µl Cas9 Protein (M0386, NEB), 0.5µl buffer (NEB), up to 5µl with ddH2O

### Tol2 Transgene

The transgene was cloned into a UAS-GFP-MiniTol2-plasmid, which was a present from the Kawakami lab (https://www.nig.ac.jp/nig/research/organization-top/organization/kawakami). The promotor was cloned using EcoNI and SalI restriction sites, the *dnm3os* lncRNA using SgrI and SnaBI.

### Zebrafish care and rearing

Zebrafish were cared and reared under standard conditions (108) and in accordance with local guidelines. Our animal database is FacileFish (109). All animal experiments are in accordance with EU and German welfare law and were approved by the local authorities of Rhineland-Palantine.

### Methylation assays

Gonads were isolated from 1 month old larvae and stored in RNAlater Solution (Invitrogen). Somatic tissue was used to assess the genotype and up to 5 gonads were pooled for DNA isolation using the NucleoSpin Tissue XS kit (Machery&Nagel) and DNA was then converted by the EZ DNA Methylation Direct kit (ZymoResearch). Analysis was done either by amplifying bi-sulfite treated DNA with the Q5U-HotStartHighFidelity Polymerase (NEB) and then analyzing it on a 3% agarose gel or by cloning amplified bi-sulfite treated DNA fragments into the Invitrogen™ Zero Blunt™ TOPO™ PCR Cloning Kit for Sequencing, with One Shot™ TOP10 Chemically Competent *E. coli* and then Sanger-sequencing the resulting clones. The results were analyzed and visualized using QUMA (110).

### RNA extraction and cDNA

RNA was extracted using Trizol (Invitrogen) together with the RNA isolation RNA Micro- or Miniprep-kits (ZymoResearch). Up to 1µg was the used for reverse transcription using the ProtoScript Reverse Transcription kit (NEB). qPCR was done using iTaq universal SYBR Green Supermix (BioRad) on a Viia7 or QuantStudio 5 (Applied BioSciences). For the miRNA-cDNA preparation we used the miRCURY LNA RT Kit (Qiuagen), together with the rno-miR214-3p LNA for the detection of miR214 and the hsa-miR-92b-3p for normalization (111).

### Microscopy and stainings

Pictures were taken on a Leica Stereo M165FC with a DFC450C camera. Photoshop (Adobe) was used to merge and assemble image panels. Bone and cartilage staining was done as described before (112).

### BioinformaMc analysis

Full name: António Miguel de Jesus Domingues

ORDCID: 0000-0002-1803-1863

### mRNA sequencing and data processing

NGS library prep was performed with Illumina’s TruSeq stranded mRNA LT Sample Prep Kit following Illumina’s standard protocol (Part # 15031047 Rev, E). Libraries were prepared with a starting amount of 193 ng and amplified in 13 PCR cycles, Libraries were profiled in a DNA 1000 Chip on a 2100 Bioanalyzer (Agilent technologies) and quantified using the Qubit dsDNA HS Assay Kit, in a Qubit 2,0 Fluorometer (Life technologies). All 16 samples were pooled in equimolar ratio and sequenced on 1 NextSeq 500 Highoutput FC, SR for 1 x 84 cycles plus 7 cycles for the index read.

The quality of sequencing reads was assessed with FastQC before being aligned against the *D. rerio* genome assembly and the genome annotation (Zv10, UCSC) with STAR v2.5.2b (– runMode alignReads –outStd SAM –outSAMattributes Standard –outSAMunmapped Within –outReadsUnmapped None –outSJfilterReads Unique –outFilterMismatchNmax 2 – outFilterMultimapNmax 10 –alignIntronMin 21 –sjdbOverhang 83). Read alignment was performed using assembled contigs.

#### Differential gene expression

Gene expression was quantified with subread featureCounts v1.5.1 (-s 2 –donotsort) (113) and differential expression analysis performed with the Bioconductor package DESeq2 v.1.18.1 (114).

Pathway enrichment was performed with the gprofiler2, via the R API for g:profiler (115, 116), using all expressed genes as background, “gSCS” correction_method, and gene sets in the following sources were tested for enrichment: “GO”, “REAC”, “MIRNA”, “CORUM”. The threshold for significant enrichment was (p-value) was set at 0.05.

### Data deposition

RAW sequencing reads and the tables for gene counts can be found in GEO with the accession number GSE254036. [reviewer token: mpsfmoqaxtyfhen]

## Acknowledgements

We thank IMB’s core facilities and our animal caretakers for excellent support, Y. ElSherif for prime technical assistance, J. B. for help with the methylation assays and F. K. for consultation on statistics. The use of IMB’s NextSeq500 funded by the Deutsche Forschungsgemeinschaft (DFG, German Research Foundation – 329045328) is gratefully acknowledged.

---

**Figure S1:**
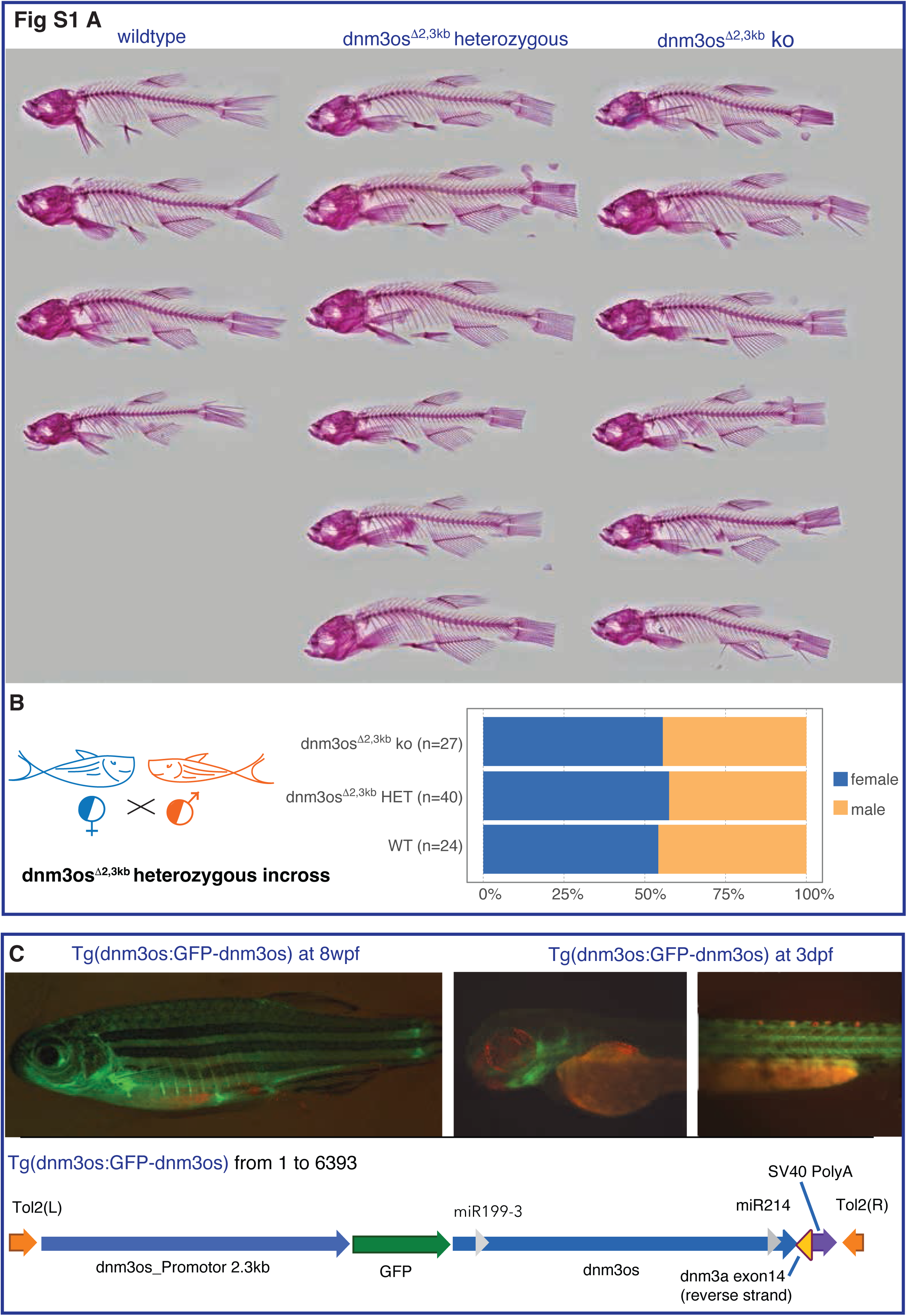
Panel **(A)** shows a bone staining of variety of adult zebrafish, in the first column wildtypes, the second column heterozygous mutants for *dnm3os^Δ2,3kb^* and in the last column homozygous *dnm3os^Δ2,3kb^* siblings. In **(B)** the sex distribution of offspring from a heterozygous incross of *dnm3os^Δ2,3kb^* fish is shown. **(C)** illustrates the transgenic line we made, a fusion of GFP with *dnm3os* driven by 2,3kb of the endogenous *dnm3os* promotor in a juvenile fish on the left side, where it mainly marks cartilage. In 3dpf old embryos on right side, we could also observe signal in the head cartilage, but also in vessels in the trunk and tail, though we did not assess, whether these vessels were blood or lymphatic vessels.

**Figure S2:**
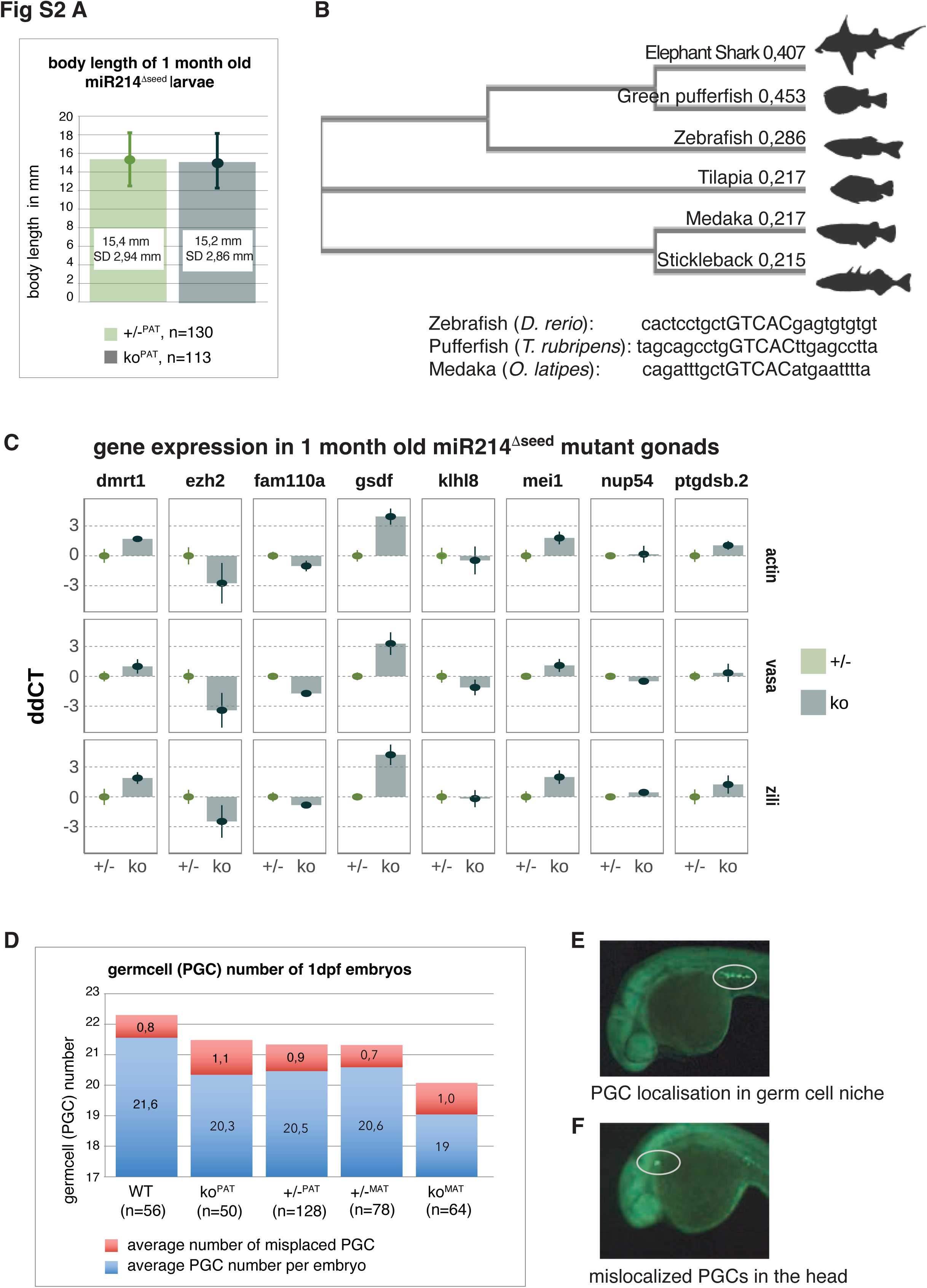
The body length of *miR214^Δseed^* larvae at 1 month were assessed in heterozygous (+/-) and homozygous mutants (ko) **(A)**. These larvae derived from a paternal cross (PAT) of homozygous *miR214^Δseed^* male to *miR214^Δseed^* heterozygous females. **(B)** Neighbour-joining tree of *gsdf* from different species using the Muscle program (117), the elephant shark was used as an outgroup. Below the conserved *miR214* binding site in the *gsdf* 3’UTR between zebrafish (*D.rerio*), Medaka (*O. latipes*) and the pufferfish (*T. rubripes*) is highlighted in capital letters. Different potential *miR214* target genes were tested in **(C)** using different reference genes: the ubiquitous β*-actin* or the germline specific genes *vasa* and *zili.* As templates served one month old heterozygous or homozygous *miR214^Δseed^* gonads, derived from a paternal cross, where homozygous *miR214^Δseed^* males were mated to *miR214^Δseed^* heterozygous females. The y-axis shows the ddCT-value (118) in an arbitrary unit. At one day post fertilization (dpf) the localization and number of <u>p</u>rimodial <u>g</u>erm <u>c</u>ells (PGCs) was assessed **(D)** using the Tg(vasa:GFP) line **(E, F)**. Embryos were either wildtype (WT), homozygous (ko) or heterozygous (+/-) for *miR214^Δseed^*. In superscript the type of cross is depicted either ‘PAT’ for male homozygous mutants to female heterozygous mutants or a reciprocal cross with homozygous females mated with heterozygous males ‘MAT’. The transgene was always conveyed via the female. In **(E)** and **(F)** are examples of PGCs, that localized properly in their niche (**E,** circled), or PGCs that mis-localized, here in the head region (**F,** circled).

**Figure S3:**
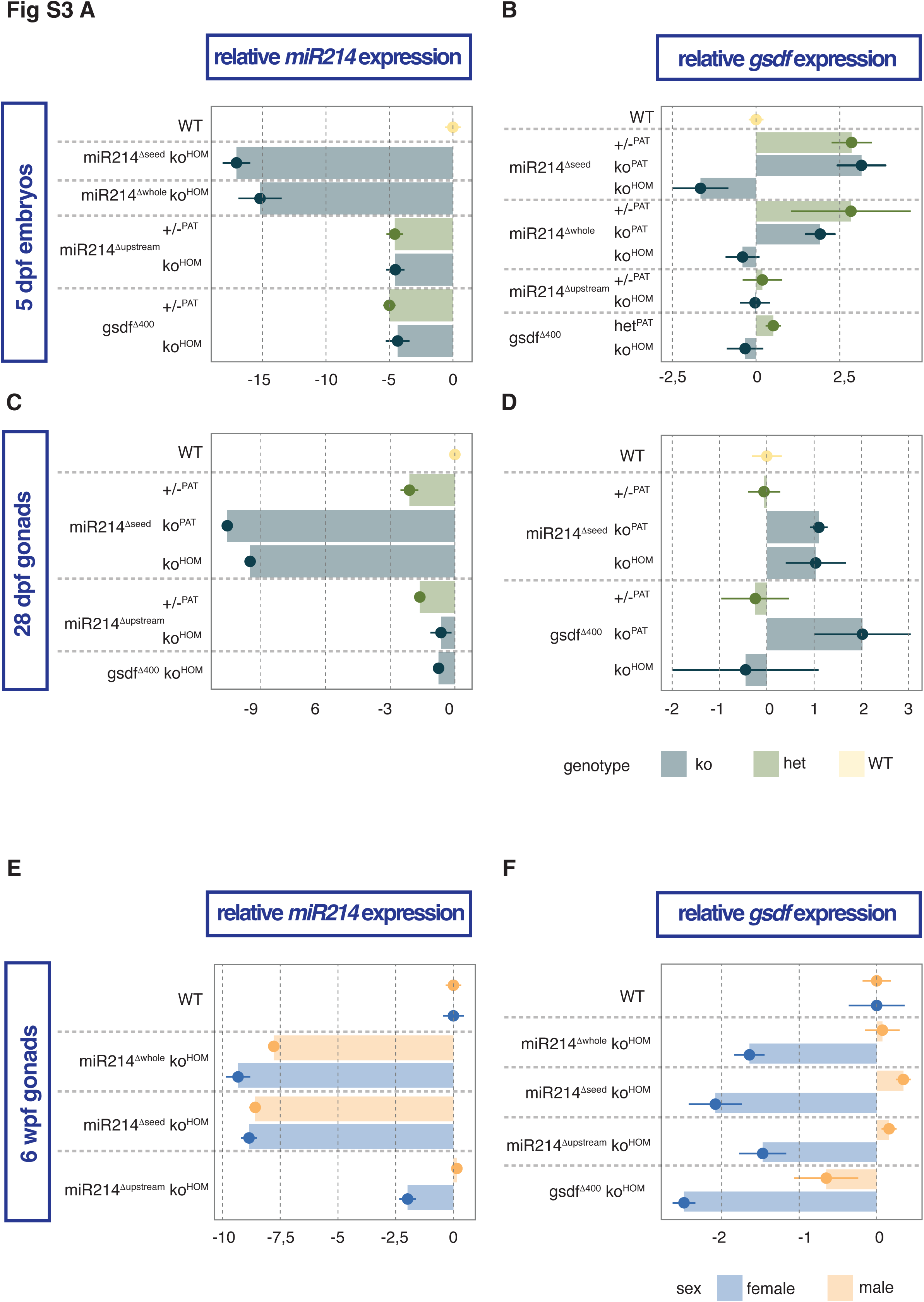
Relative *miR214* (**A, C, E)** or *gsdf* (**B, D, F)** expression in whole 5dpf embryos (**A, B)**, 28dpf gonads (**C, D)** or 6-week-old gonads (wpf) (**E, F).** The x-axis shows the relative RNA expression as ddCT (118) in an arbitrary unit, β*-actin* was used as a reference gene. The genotype and the allele are depicted on the y-axis. The superscripted letters describe the type of cross the analyzed fish were derived from: ‘HOM’ = parents were both homozygous carrier for describes mutations, ‘PAT’ = these larvae derived from a cross between a heterozygous female with a homozygous male. ‘WT’ = wildtype. The gonads at 6wpf in **E** and **F** were already differentiating into ovary and testis and were hence divided into male and female.

**Figure S4:**
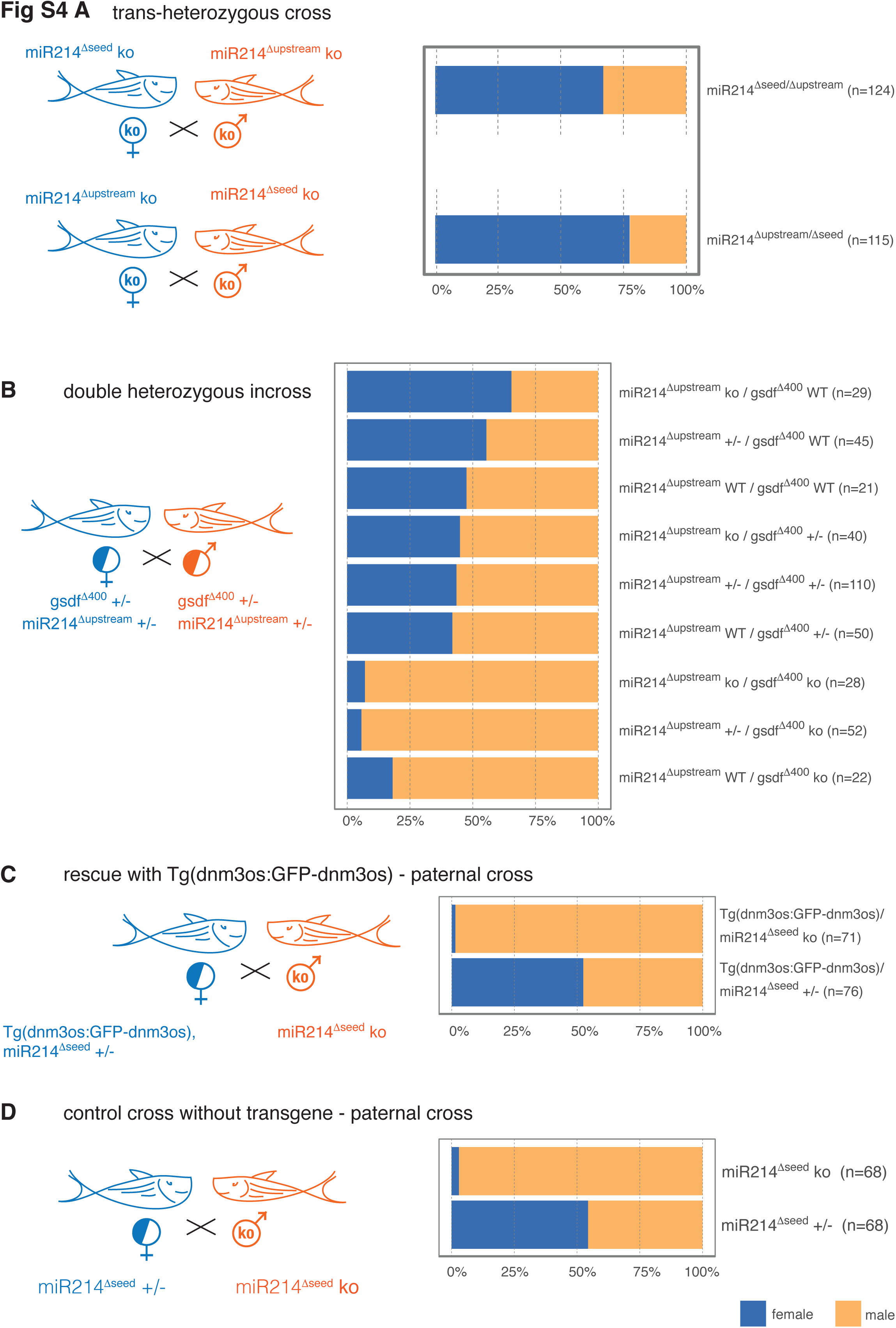
Homozygous *miR214^Δseed^* or *miR214^Δupstream^* animals were mated in reciprocal crosses and their trans-heterozygous offspring was examined for their sex distribution **(A)**. Double heterozygous animals for *gsdf^Δ400bp^* and *miR214^Δupstream^* were crossed and the sex distribution of the offspring was analyzed according to genotype **(B).** We then tried to rescue the *miR214^Δseed^* male bias, by rescuing it with the *Tg(dnm3os:GFP-dnm3os)* **(C)**, here the transgene was conveyed via the females, which were also heterozygous for the *miR214^Δseed^* allele and crossed to homozygous *miR214^Δseed^* males. As a control we performed the same cross, but without the transgene **(D)**.

**Figure S5:**
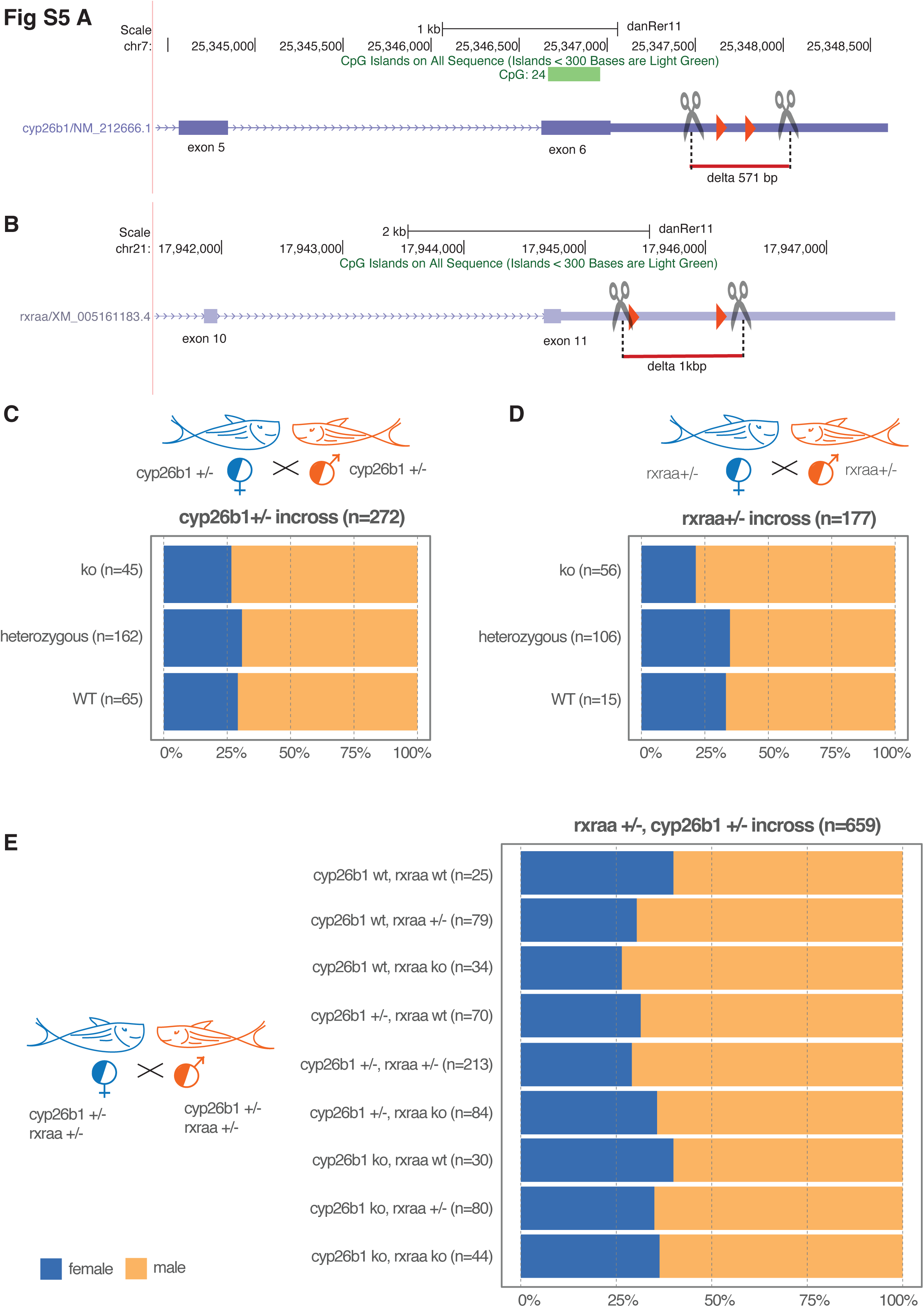
Gene structure of *cyp26b1* **(A)** and *rxraa* **(B)** showing the position of the potential *miR214* binding side as red triangles and the position of the sgRNA used for mutagenesis as scissor pairs. We then used those mutants and incrossed heterozygous carriers to analyze the sex distribution of their offspring in **(C)** and **(D).** We also mated double heterozygous carrier fish for *rxraa* ^+/-^ and *cyp26b1* ^+/-^ and examined the sex distribution of their offspring in **(E).** All these crosses were done once, and ‘n’ represents the total number of animals.

**Figure S6:**
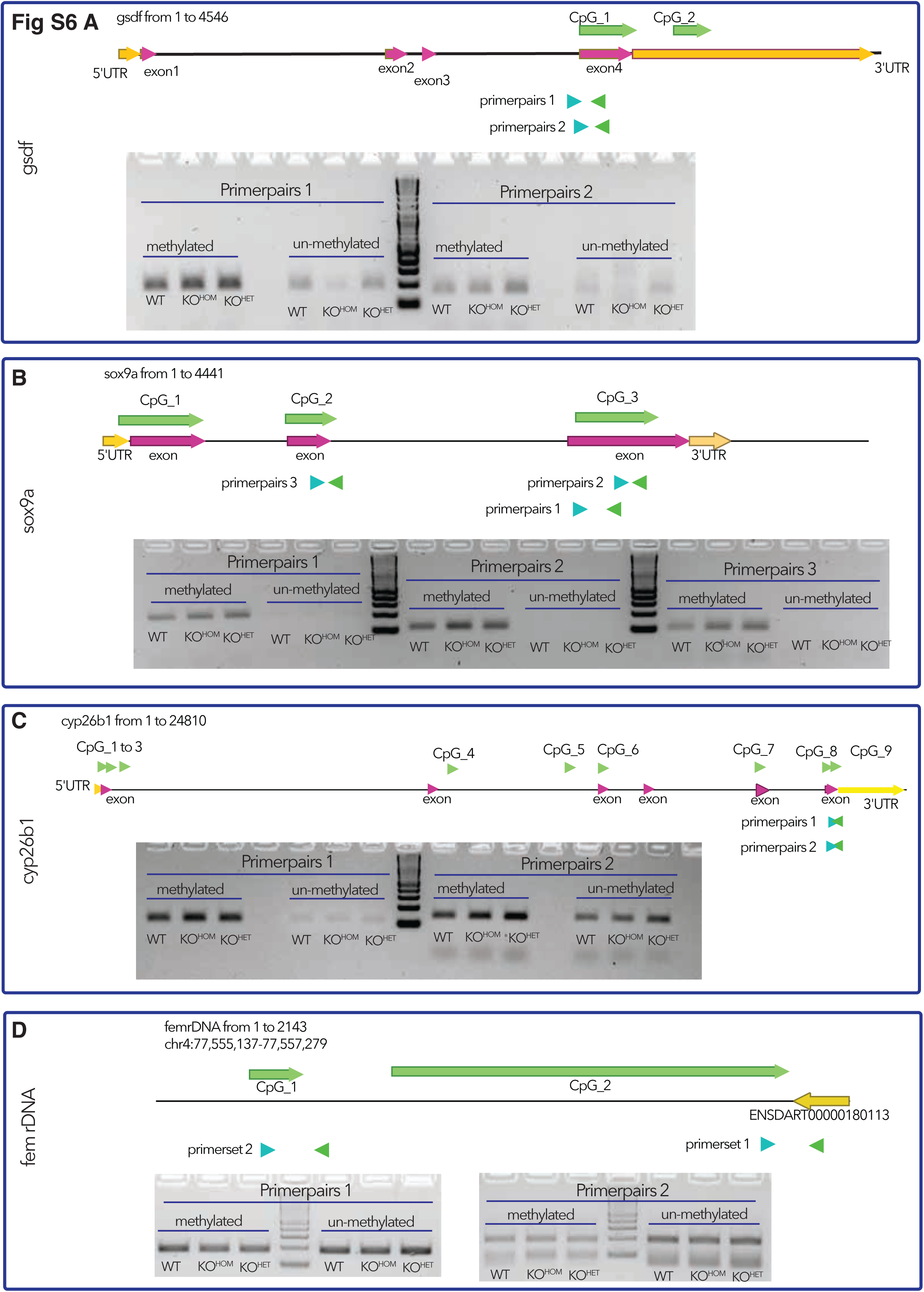
Assessment of the methylation status of different genes using methylation specific PCR (119). On top of each panel the gene structure, the CpG-Islands and used PCR-Primer are depicted. Below are the resulting agarose gels, on the left-side methylation-specific Primers, and on the right-side primer to amplify un-methylated parts were used (119). In the middle is a 1kb+ DNA-marker (NEB). Template were 1 month old gonads from wildtype, homozygous or heterozygous *miR214^Δseed^* fish. **(A)** shows the analysis for *gsdf*, **(B)** for *sox9a*, **(C)** for *cyp26b1* and for *femrDNA* **(D)**.

**Figure S7:**
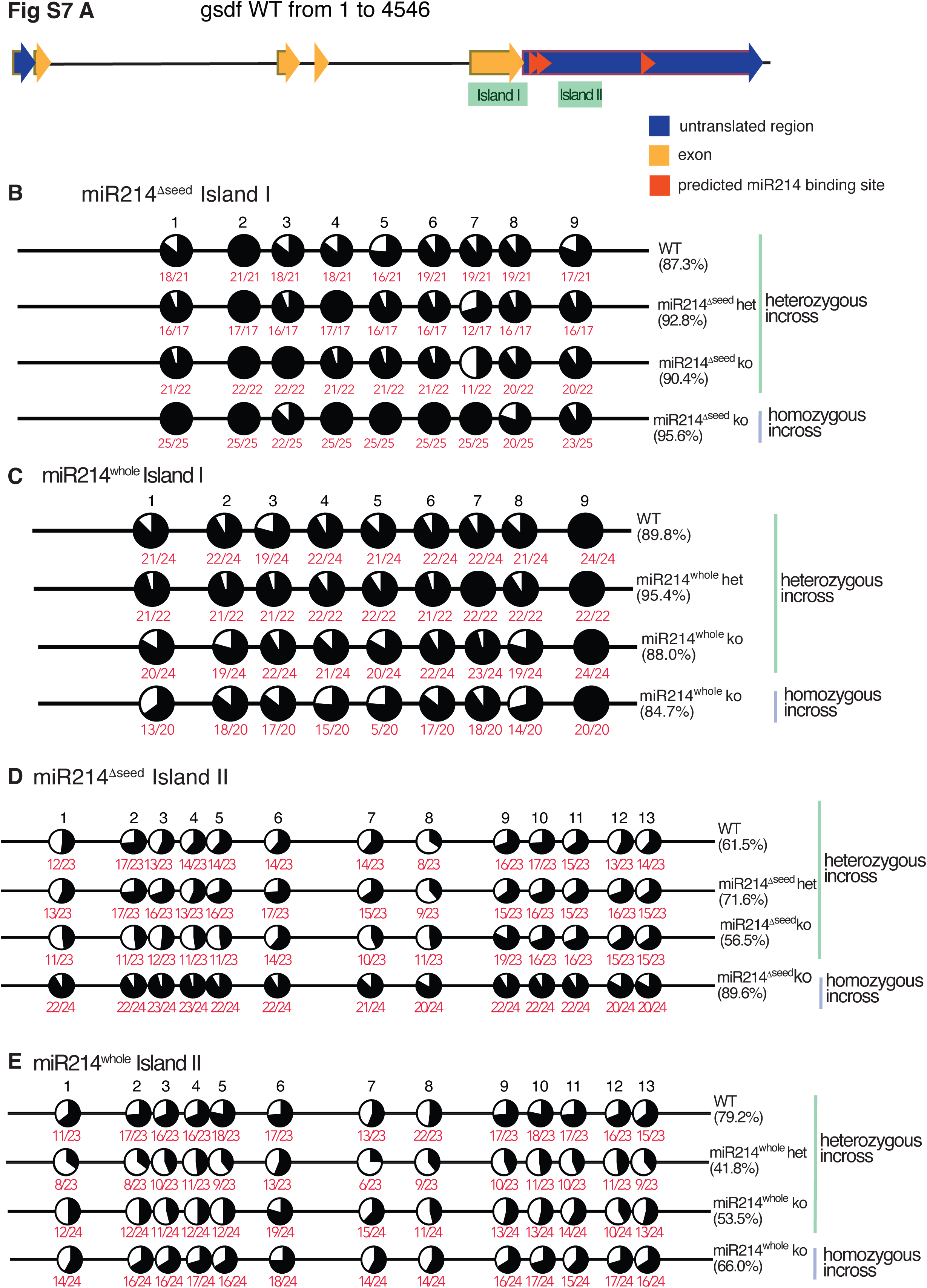
Gene structure of *gsdf*, highlighting the assessed CpG-Islands I and II **(A).** The methylation status of 1 month old gonads was analyzed using colony-sequencing (120) either by amplification of CpG-Island I **(B, C)** or CpG-Island II **(D, E)**. Gonads were derived from *miR214^Δseed^* **(B, D)** or *miR214^Δwhole^* 1 month old larvae **(C, E)**. The genotype is depicted on the right side of each lollipop diagramm. On the right-hand site, the genotype and the type of cross are indicated. The amount of black in the circles indicate the amount of methylated Cs, the more the lollipop is filled, the higher the methylation rate. The red numbers underneath the circles show the actual number of converted colonies compared to the amount of totally sequenced colonies.

**Figure S8:**
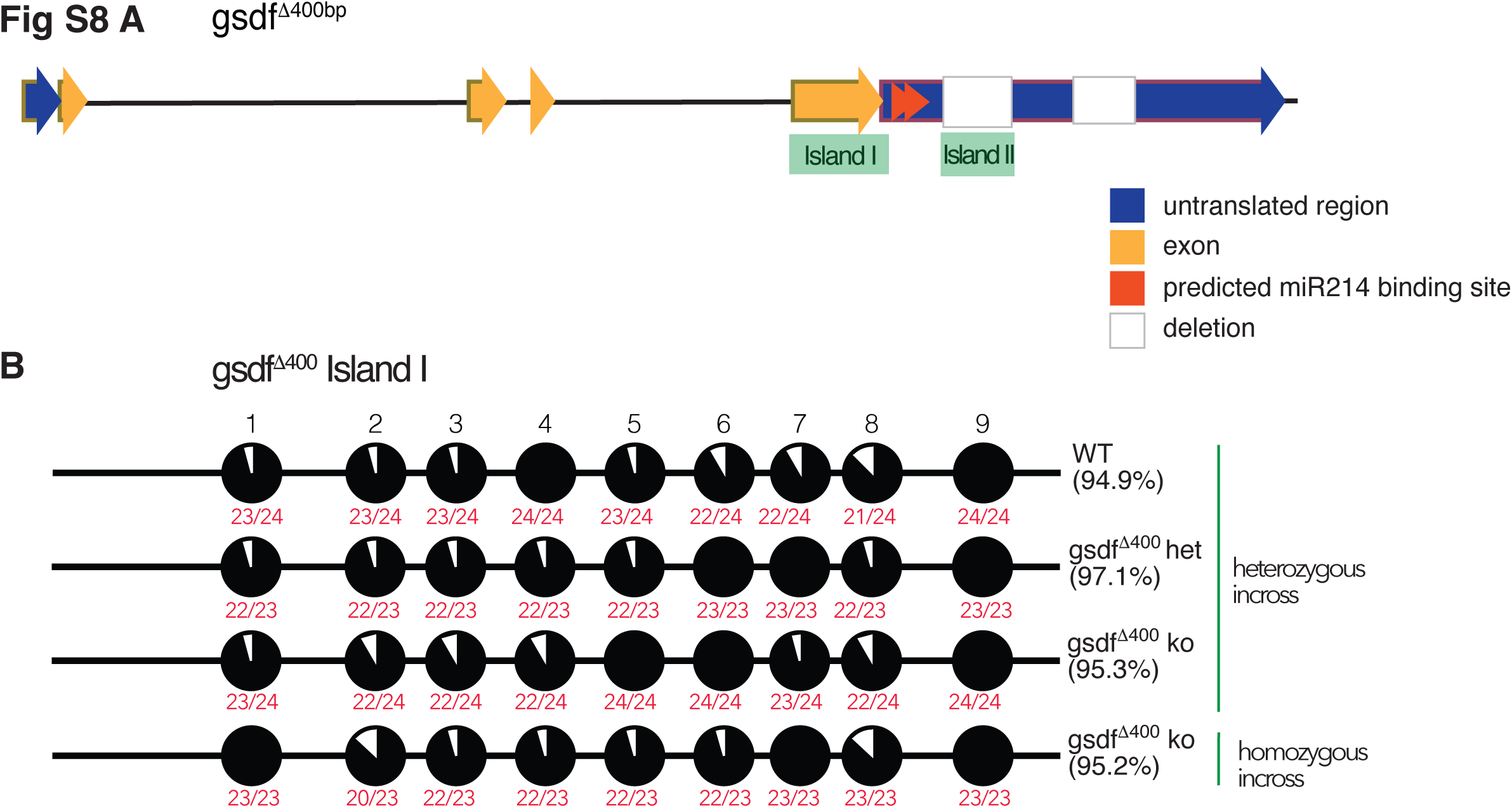
Methylation status analysis of the *gsdf^Δ400bp^* mutation **(A)** using colony-PCR and sequencing (120) for Island I **(B)** as CpG-Island 2 is deleted. Percentages given are the amount of methylation within the colonies analyzed for the respective CpG Island and numbers in red give the number of methylated colonies over the total number of colonies sequenced.

**Supplemental Table 1:**
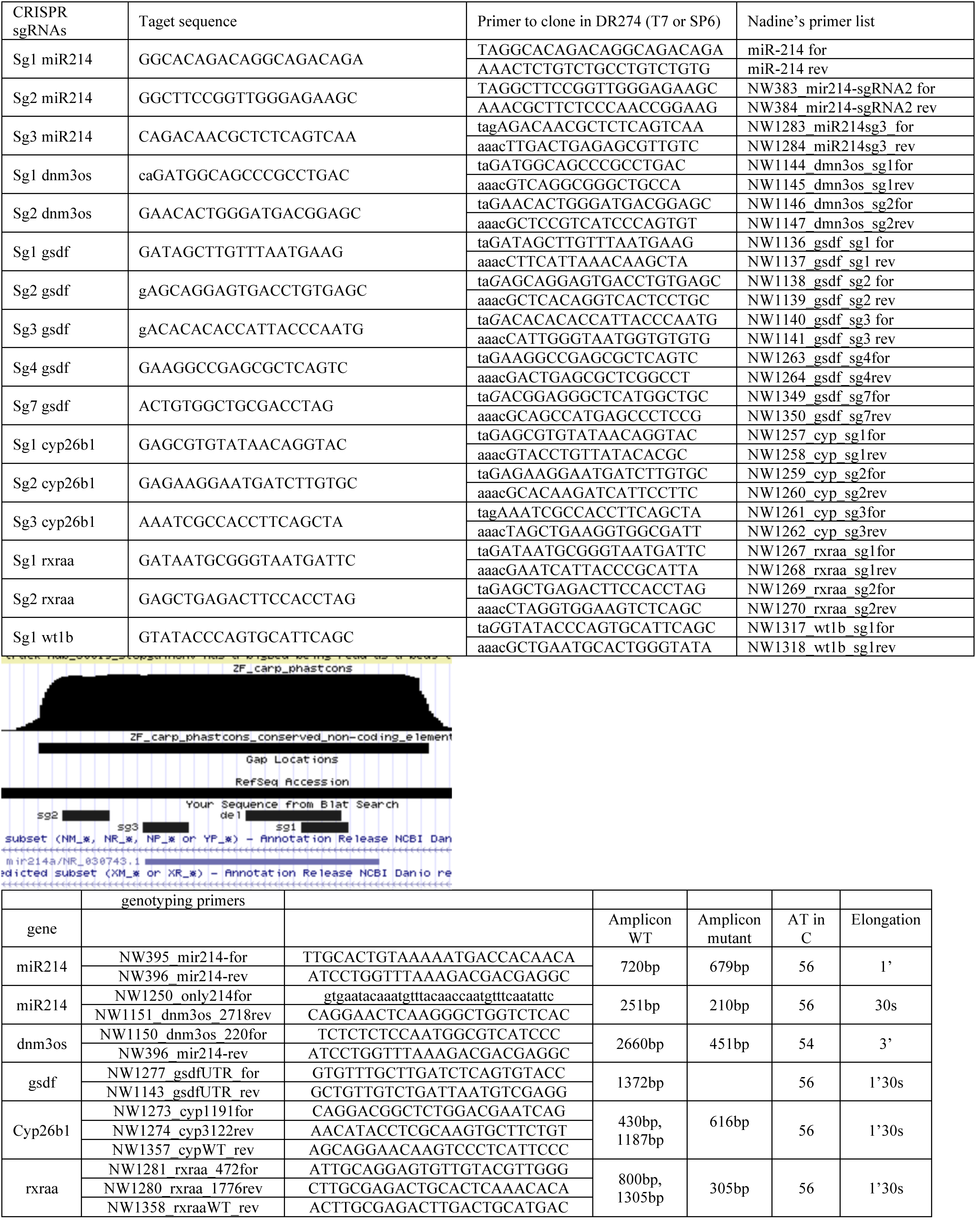
PaternalKOvsPaternalHET.annotated.xls Comparison of gene expression in 1 month old gonads between larvae obtained from a cross between homozygous *miR214^Δseed^* males to heterozygous *miR214^Δseed^* females (paternal cross).

**Supplemental Table 2:**
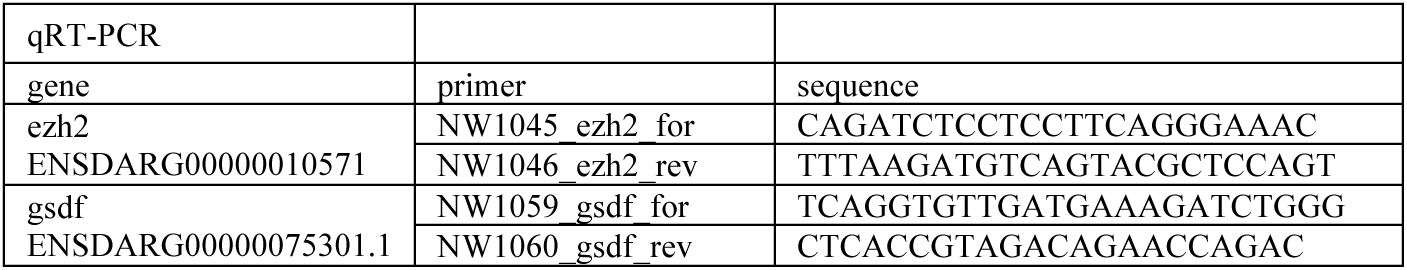
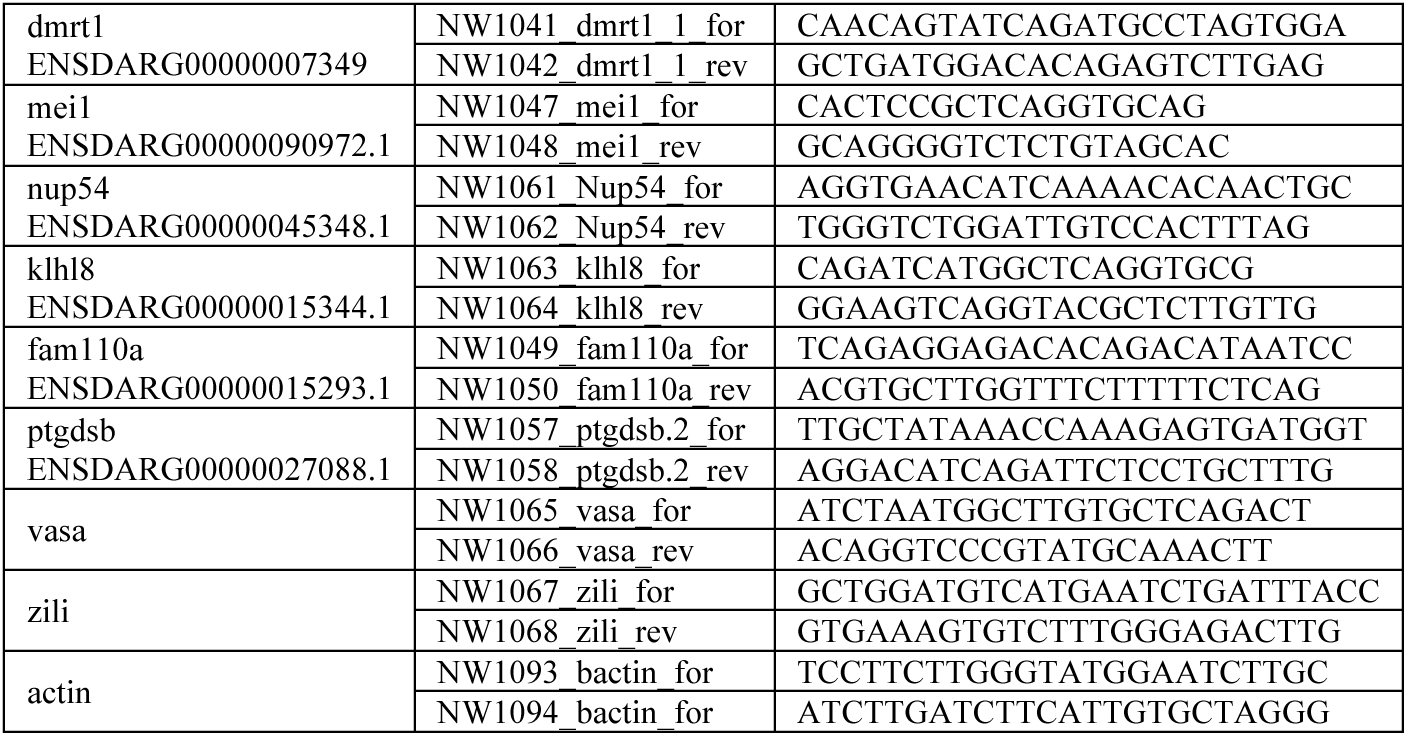
maternalKOvsmaternalHET.annotated.xls Comparison of gene expression in 1 month old gonads between larvae obtained from a cross between homozygous *miR214^Δseed^* males to heterozygous *miR214^Δseed^* females (maternal cross).

**Supplemental Table S3:**
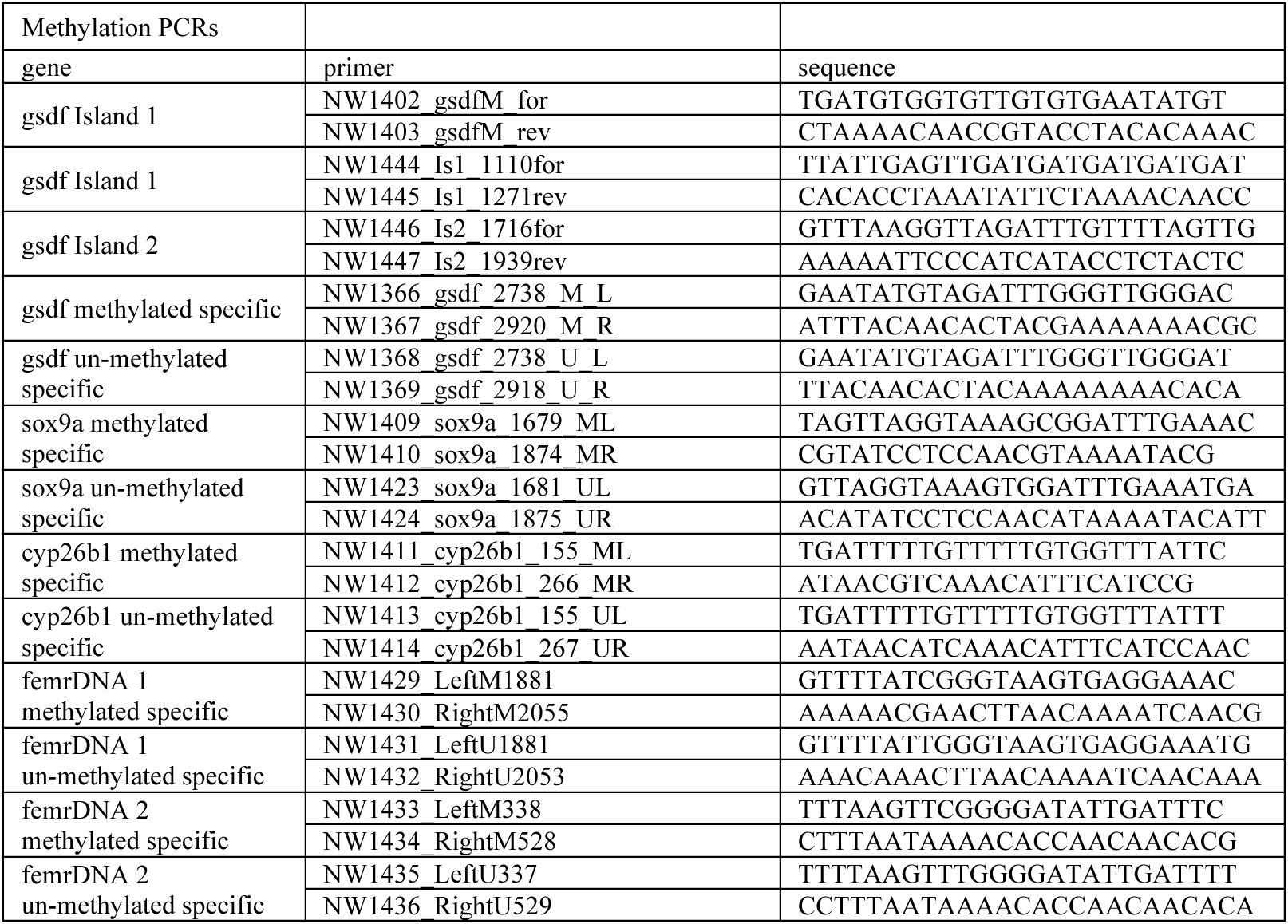
PaternalKOvsmaternalKO.annotated.xls. Expression comparison between the paternal and maternal crosses

**Supplemental Table S4:**
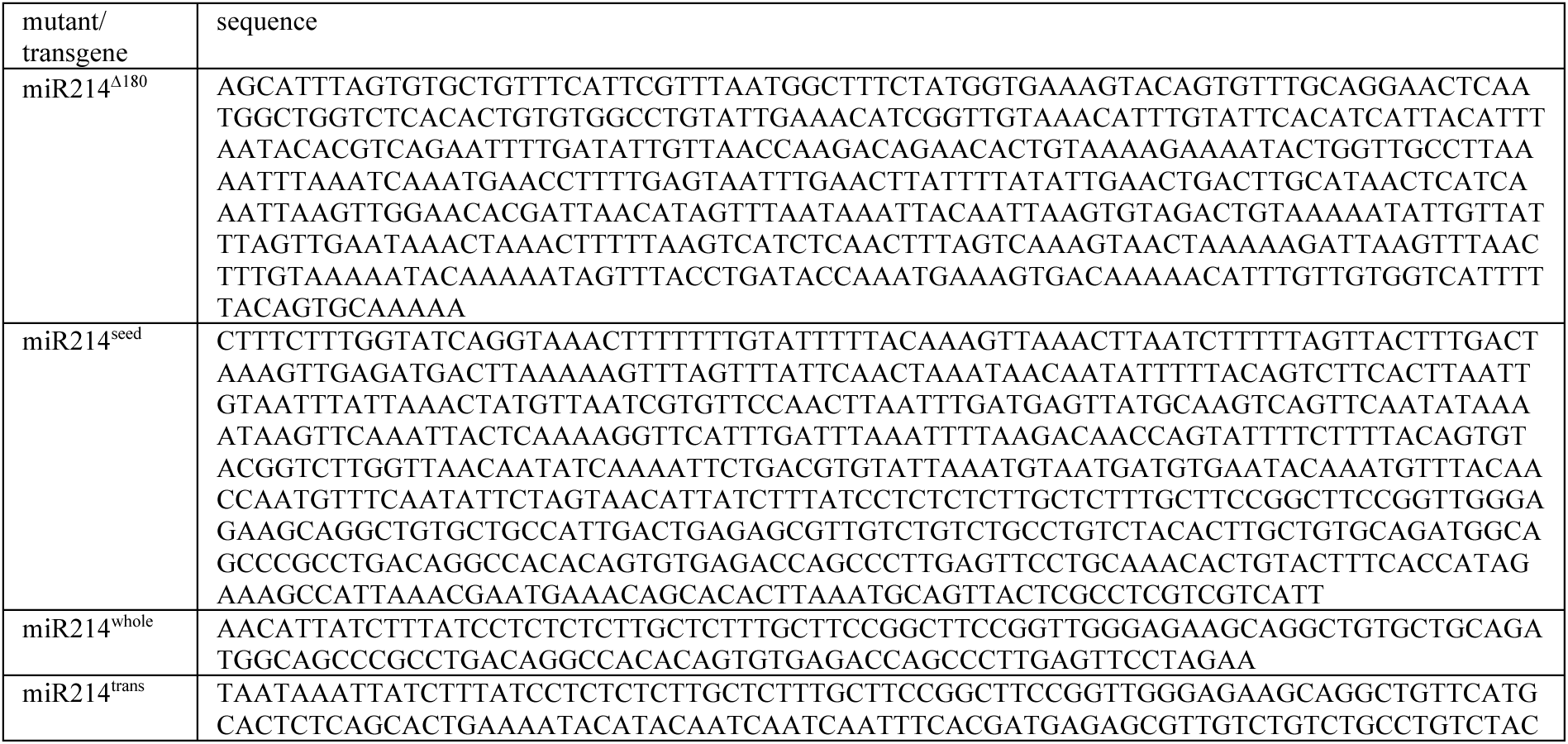
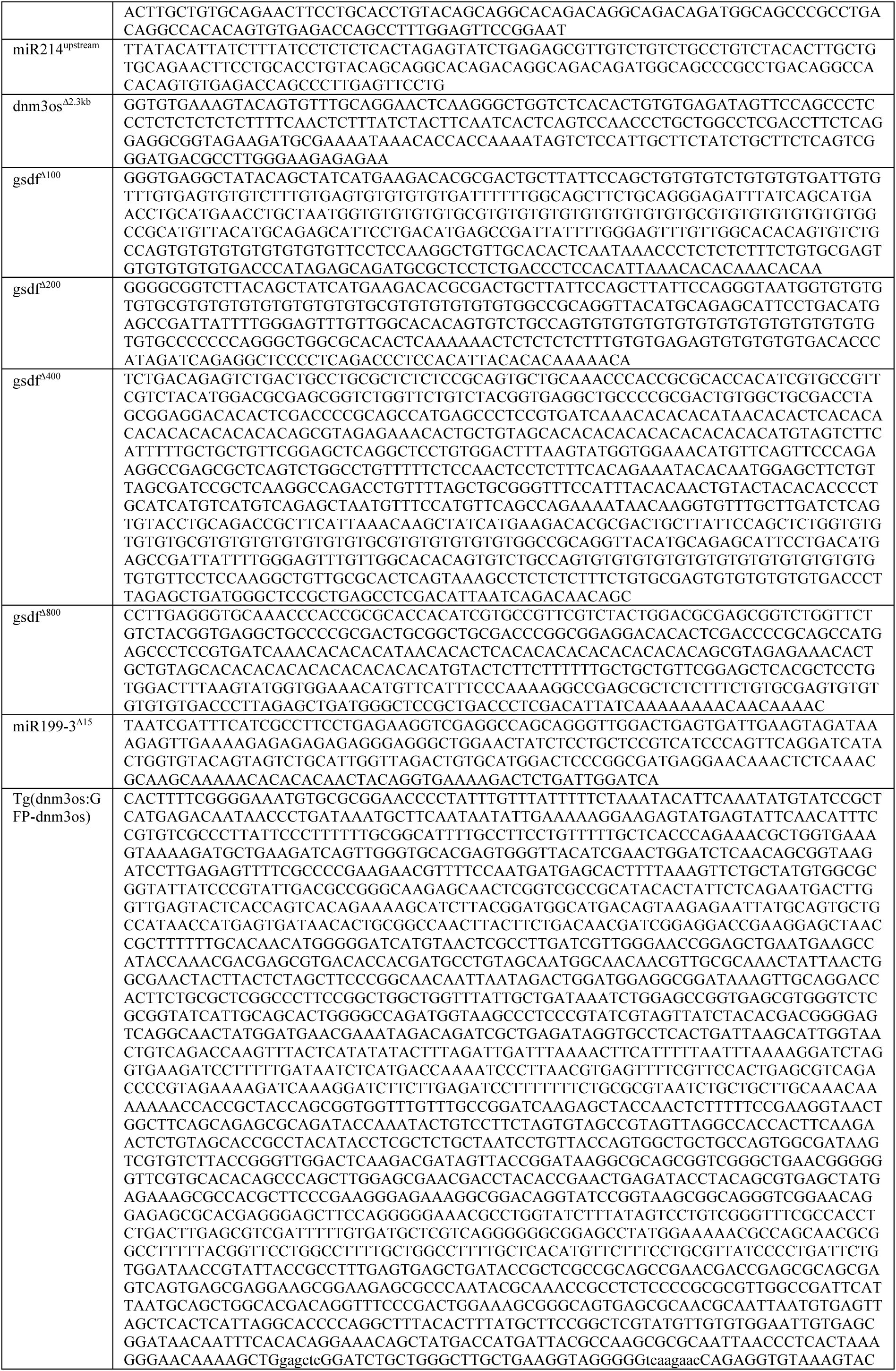
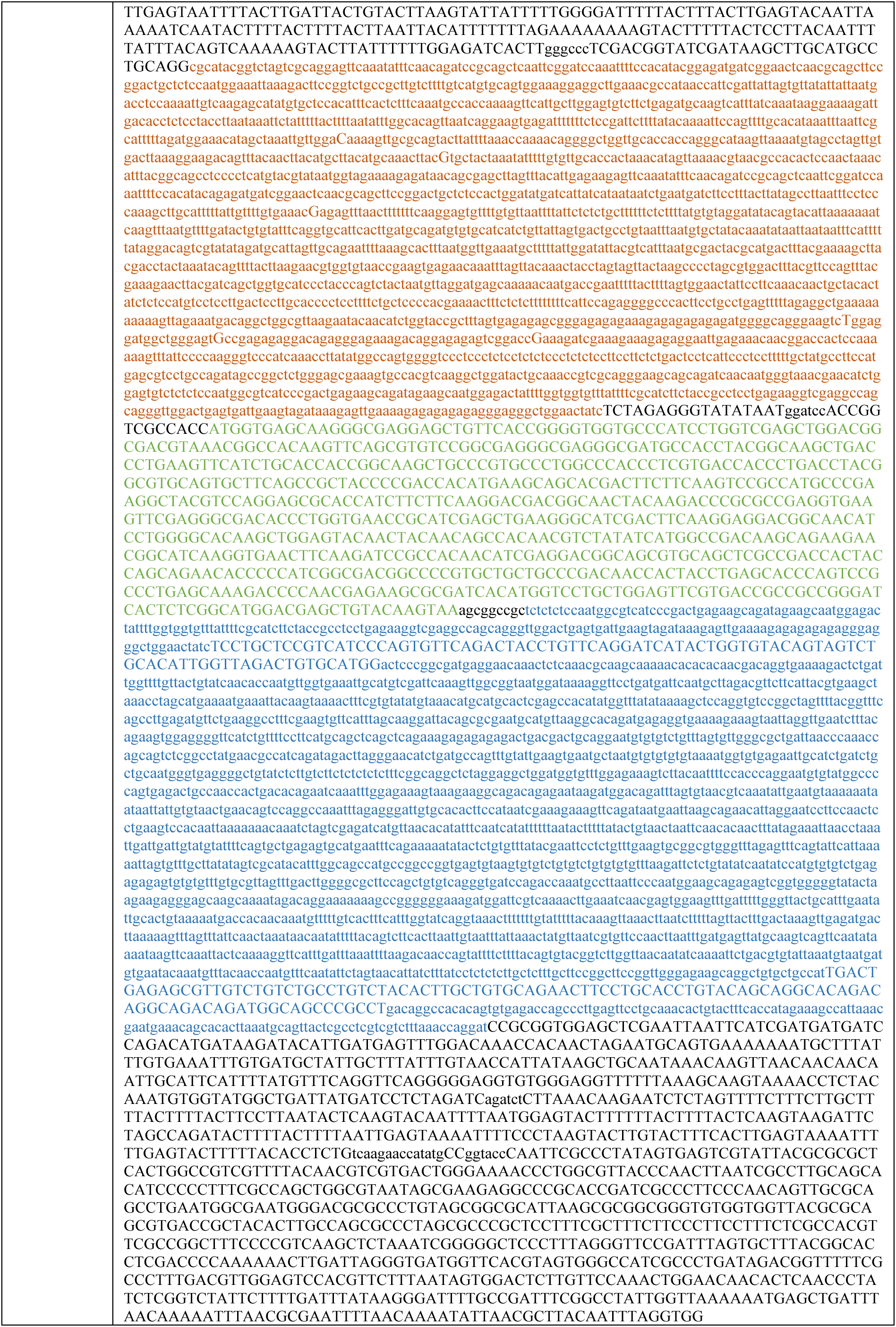
List of used oligos and sequences of generated mutant and transgenic lines.

## Notes

### Competing Interest Statement

The authors have declared no competing interest.

